# Discovery of a Cell-Permeable Gi-Signaling Inhibitor

**DOI:** 10.64898/2026.08.02.742274

**Authors:** Amira Naimi, Tim Berger, Philipp Dahlhaus, Marcel Rossol, Jonas Mühle, Wieland Steinchen, Gert Bange, Uwe Linne, Paolo Stincone, Xavier Deupi, Gebhard Schertler, Lars Jürgenliemke, Evi Kostenis, Jens Kockskämper, Daniel Hilger, Daniel Petras, Raphael Reher

**Author notes:** These authors contributed equally (Amira Naimi, Tim Berger).

## Abstract

Heterotrimeric G proteins regulate diverse physiological processes, yet selective small molecule modulators for the Gα_i_ subfamily remain scarce. Here, we employed native metabolomics to screen a fungal culture collection for G protein binders, which led to the discovery of *N*-hydroxyapiosporamide (NHAP), a fungal specialized metabolite that functionally inhibits Gi-signaling. NHAP selectively binds Gα_i1_ and diminishes Gα_i1_-mediated GTP turnover in biochemical assays. In primary ventricular cardiomyocytes, NHAP largely reversed acetylcholine-induced negative inotropy, demonstrating functional Gi-signaling blockade in a physiological context without acute cytotoxicity. This work establishes NHAP as a first-in-class Gi-selective inhibitor, providing a cell-permeable chemical scaffold for the optimization of next-generation agents to control Gi-signaling in cell-based systems and, ultimately, to target Gα_i1_-driven pathologies.

## Introduction

Heterotrimeric G proteins serve as central transducers in G protein-coupled receptor (GPCR) signaling cascades, controlling numerous critical processes in human physiology.^1–3^ They comprise three polypeptide chains: a Gα subunit and an obligate Gβγ heterodimer.^4^ The Gα subunit functions as a molecular switch, cycling between inactive and active states depending on the bound guanosine nucleotide.^5^ In the inactive state, Gα is bound to guanosine diphosphate (GDP) and associated with Gβγ. Activation occurs upon receptor-mediated exchange of GDP for guanosine triphosphate (GTP), inducing conformational changes that lead to Gβγ dissociation and the regulation of downstream effectors. Signaling is terminated by the intrinsic GTPase activity of the Gα subunit, which hydrolyzes GTP to GDP, resulting in reassociation with Gβγ.^6^ In humans, 21 distinct Gα subunits are categorized into four subfamilies: Gα_q/11_, Gα_s_, Gα_i/o_, and Gα_12/13_^.7–9^ Consistent with their vital roles in cellular homeostasis, dysregulation of G protein activity is associated with various diseases, including uveal melanoma, McCune-Albright syndrome, and endocrinopathies.^10–13^ Despite their clinical relevance, direct G protein-targeting agents are severely underrepresented in the current pharmacopoeia.^14–16^ In stark contrast, GPCRs represent the most thoroughly investigated drug targets. With over 800 identified human GPCRs, it is unsurprising that more than one-third of all FDA-approved drugs target GPCRs, primarily as selective agonists or antagonists.^17–19^ Direct targeting of heterotrimeric G proteins, however, has proven far more challenging. They have long been considered “undruggable” due to their intracellular location, high affinity for endogenous nucleotides, and high structural conservation across different Gα subfamilies.^14,20–22^ To date, only a few selective small molecule inhibitors of G protein signaling have been identified. For the Gα_q/11/14_ subfamily, the natural products FR900359 (FR) and YM-254890 (YM) are widely used as pharmacological tools.^23–26^ While FR initially has been isolated from an endosymbiotic plant-bacterium system, both, FR and YM have also been described to be produced by free-living soil bacteria from the genus *Chromobacterium*. These cyclic depsipeptides act as guanine nucleotide dissociation inhibitors (GDIs) that additionally stabilize the inactive heterotrimer.^27–29^ However, attempts to develop inhibitors for other G protein families based on the FR scaffold have been unsuccessful, suggesting that co-evolution of the biosynthetic gene cluster of FR and Gα_q_ led to remarkable selectivity that is difficult to engineer synthetically.^25,30^ For the Gα_s_ subfamily, macrocyclic peptides (GD20, GN13) were recently discovered, but they suffer from limited cellular activity.^31^ For the Gα_i_ family, Pertussis Toxin (PTX) from *Bordetella pertussis* is the only widely used inhibitor.^32^ While valuable in foundational research, PTX is a protein toxin that acts irreversibly and requires endocytosis for intracellular trafficking, leading to slow and variable effects. There is currently no small molecule inhibitor that offers reversible, tunable, and isoform-selective inhibition of Gi-signaling for precise temporal control *in vitro* and *in vivo*. Macrocyclic peptides have been explored to fill this gap, with candidates such as cycGiBP, KB-752, and GPM-1-3 showing specific binding to Gα_i1_^.33–37^ However, most require extensive chemical modifications to achieve cellular permeability, limiting their utility as tool compounds to study cellular processes. This underscores the need for new chemical scaffolds. Natural products, particularly from endosymbiotic systems, pose fruitful sources for discovery of cell permeable G protein modulators, as demonstrated by the obligate plant-bacterium endosymbiosis responsible for FR production. Marine endosymbionts, such as those associated with tunicates, are speculated sources of bioactive metabolites (e.g., sameuramide A), but no marine small molecule directly modulating Gi-signaling has been identified so far.^38–40^

To address this, we hypothesized that marine-derived fungal endosymbionts harbor unique G protein modulators. In fungi, Group I Gα proteins share significant sequence homology to mammalian Gα_i1_, including conserved myristoylation and ADP-ribosylation motifs.^39,41,42^ Traditional discovery pipelines are redundant and time-consuming, separating isolation and structural annotation from bioactivity determination. To overcome this, we have developed a scalable native metabolomics approach that integrates non-targeted LC-MS/MS with protein binding detection via native mass spectrometry.^43–45^ This method enables bioactivity-focused compound identification from complex mixtures without prior separation.

In this study, we report the discovery of *N*-hydroxyapiosporamide (NHAP) as a cell-permeable small molecule Gi-signaling inhibitor, identified using native metabolomics. Biochemical and pharmacological evaluation reveals that NHAP functionally inhibits Gi-mediated signaling in primary cardiomyocytes, while its structural analog apiosporamide (AP) shows no specific activity in the latter. This work provides the first selective small molecule pharmacological tool for the optimization of next-generation agents to probe Gi signaling in research and to target Gi-driven pathologies.

## Results

### Native metabolomics platform enables discovery of Gα_i1_ modulators

To identify direct G protein modulators from complex natural product extracts, we employed a native metabolomics workflow integrating non-targeted LC-MS/MS with protein binding detection via native mass spectrometry (native MS).^43–45^ Method optimization ensured nativelike conditions for the purified Gα_i1_ protein (20 µM), yielding stable signals and minimizing nonspecific binding (**Fig. S1**). To validate the platform, we screened the selective Gα_q/11/14_ inhibitor FR against representative members of the Gα_i_, Gα_s_, and Gα_q_ families. As expected, FR bound specifically to Gα_11_ and an engineered FR-sensitive Gα_i1_ mutant, but not to wildtype Gα_i1_ or Gα_s_ (**Fig. S2**), confirming the system’s ability to distinguish selective binders.

### Discovery of selective Gα_i1_ binders from marine fungal extracts

We screened 40 extracts from 10 endosymbiotic marine fungi against Gα_i1_. Two extracts contained features showing specific mass shifts indicative of protein binding. From *Apiospora* sp. 589, we identified two co-occurring features at *m/z* 430.2218 and *m/z* 446.2160 (**Fig. 1a–d, S3**). A third binder from *Stachylidium bicolor* 293 K04 was identified as the known peptide endolide E (**Fig. S4**).^46^ To assess specificity, we isolated the features from *Apiospora* sp. 589 and subjected them to selectivity profiling against Gα_i1_, Gα_s_, Gα_11_, and Gβγ subunits via native MS. The feature at *m/z* 446 bound exclusively to Gα_i1_ in its native-like state, with no binding observed to denatured Gα_i1_ protein or other G protein families (**Fig. 1e–h**). In contrast, the feature at *m/z* 430 displayed unselective binding to all tested Gα subunits and the Gβγ dimer (**Fig. 1i–l**). Endolide E similarly showed strong, but non-specific binding (**Fig. S4**). Based on this selectivity profile, we prioritized *m/z* 446 for functional characterization, while retaining *m/z* 430 as a putative structural analog for initial structure-activity studies.

**Fig. 1:**
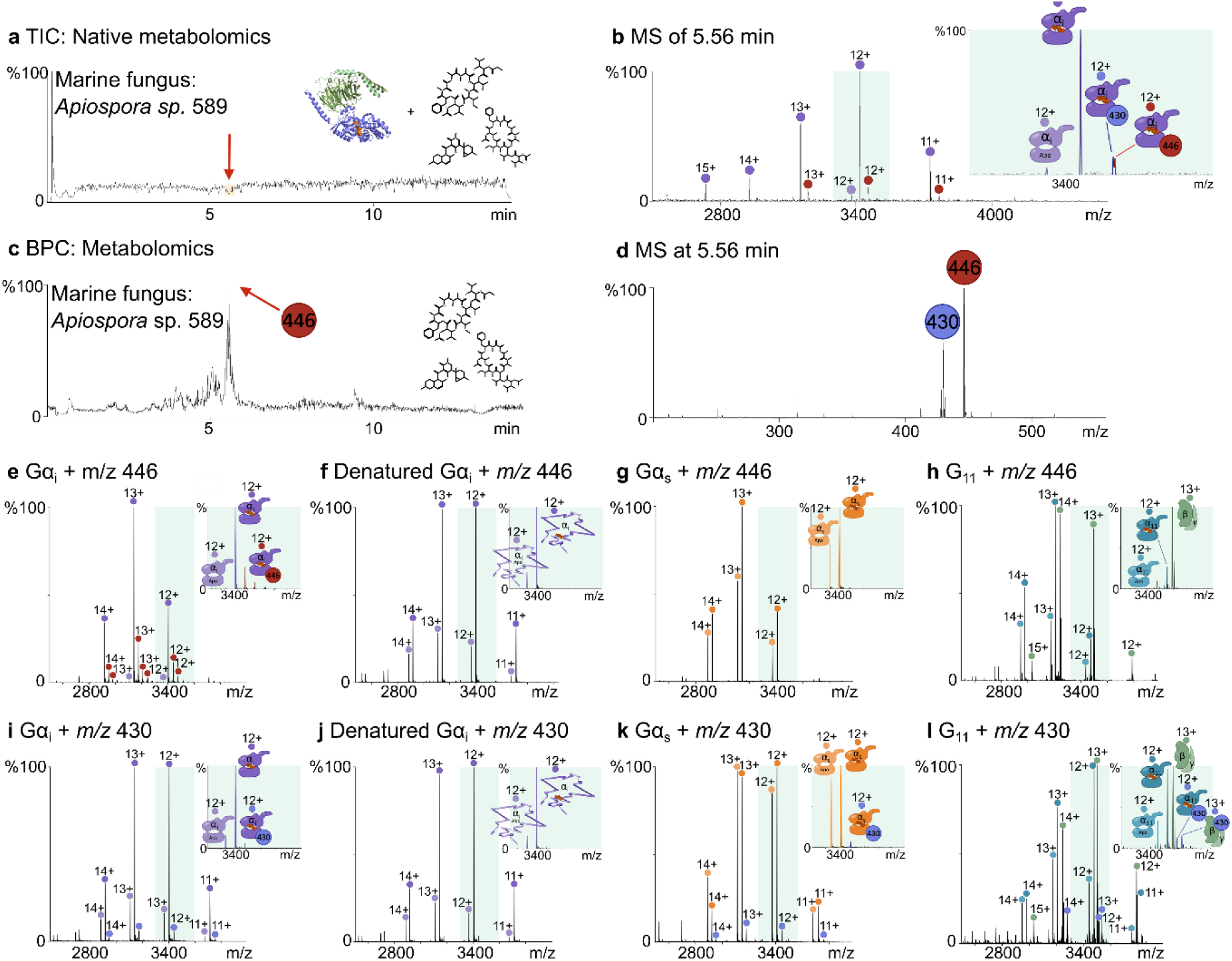
Native metabolomics-guided identification of Gα_i1_ binders. **a** Total ion chromatogram (TIC) of the native metabolomics run with Gα_i1_ and *Apiospora* sp. 589 extract. **b** Native mass spectrum at 5.56 min. Apo-Gα_i1_ (light purple) and GDP-Gα_i1_ (dark purple) signals shown. Red signal indicates GDPGα_i1_ bound to ligand (*m/z* 446) and light purple signal indicates GDP-Gα_i1_ bound to ligand (*m/z* 430). **c** Base peak chromatogram (BPC) of extract metabolomics run showing dominant peak at 5.56 min (*m/z* 446.2160). **d** HR-MS1 spectra at 5.56 min. **e-h** Selectivity tests for *m/z* 446 (NHAP) and *m/z* 430 (AP). **e** Native MS spectrum shows binding of *m/z* 446 (10x excess) to native Gα_i1_. **f** Native mass spectra of denatured Gα_i1_ and GDP-Gα_i_ (charge states in 12+). No binding of *m/z* 446. **g** Native mass spectra of Apo-Gα_s_ in light orange and GDP-Gα_s_ in orange (charge states in 12+ to 14+) and potential binder *m/z* 446 (10x excess, no mass shift). **h** Native mass spectra of heterotrimeric Gα_11_βγ (charge states of GDPGα_11_ in light blue and Apo-Gα_11_ in blue in 12+ to 15+ and Gβγ in 12+ to 15+ in green) and the potential binder *m/z* 446 (10x excess, no mass shift to neither α-subunit, nor βγ-subunit. **i** Native mass spectra of Gα_i1_ and *m/z* 430 (light purple, 10x excess). Mass shift of GDP-Gα_i_*m/z* 430 in purple. **j** Native mass spectra of denatured Gα_i1_ and GDP-Gα_i1_ (charge states in 12+, no binding of *m/z* 430). **k** Native mass spectra of Gα_s_ and *m/z* 430 (10x excess, mass shift in purple). **l** Native mass spectra of G_11_ (charge states of GDP-Gα_11_ in light blue and Apo-Gα_11_ in blue in 11+ to 15+ and Gβγ in 12+ to 15+ in green) and *m/z* 430 (5x excess, purple, charge states of Gβγ*m/z* 430 in 12+ to 14+, mass shift in purple to the α-subunit and purple to the βγ-subunit).

### Targeted isolation and structure elucidation of two structurally related Gα_i1_ binders

To elucidate the structures of the Gα_i1_ binders, we first conducted a large-scale cultivation and purification/ isolation via size-exclusion chromatography and semi-preparative HPLC, guided by LCMS-based-time course experiments to determine the best time point to harvest the producing fungi to obtain sufficient amounts of the targeted molecules (**Fig. 2a, S5**). The employed computational metabolomics (MS/MS and NMR-based) workflow accelerated this process. SIRIUS framework analysis^47^ predicted molecular formulas of C_24_H_32_NO_6_^+^ [M+H]^+^ (*m/z* 430.2218, *m/z* feature 430) and C_24_H_32_NO_7_^+^ [M+H]^+^ (*m/z* 446.2160, *m/z* feature 446), suggesting a mass difference of one oxygen atom. CANOPUS^48^ and CSI:FingerID^49^ searches proposed 4-hydroxy-2-pyridone substructures, annotating the compounds as apiosporamide (*m/z* feature 430) and *N*-hydroxyapiosporamide (*m/z* feature 446), respectively (**Figs. S6, S7**). GNPS molecular networking^50^ confirmed their structural similarity (**Fig. 2b**). Complementary, NMR data for these compounds were obtained followed semi-automated analysis of ^1^H-^13^CHSQC NMR spectra with the deep learning-based NMR tools SMART^51,52^ and DeepSAT,^53^ allowing for dereplication of natural products and compound classification. This led to the assignments of two very similar 4-hydroxy-2-pyridone-containing natural products (**Fig. 2b, 2c, S8-11, Tab. S1-2**). Bringing together the structure assumptions from the SIRIUS framework and SMART with manual analysis of the complete sets of 1D/2D NMR data (**Fig. S8-22, Tab. S3-4**) the planar structures were finally determined to be apiosporamide (*m/z* feature 430) and *N*-hydroxyapiosporamide (*m/z* feature 446), respectively. Comparison of optical rotation values with literature data identified the compounds as (−)-apiosporamide (AP, *m/z* feature 430) and (−)-*N*-hydroxyapiosporamide (NHAP, *m/z* feature 446) (**Fig. 2d**).^54–57^

**Fig. 2:**
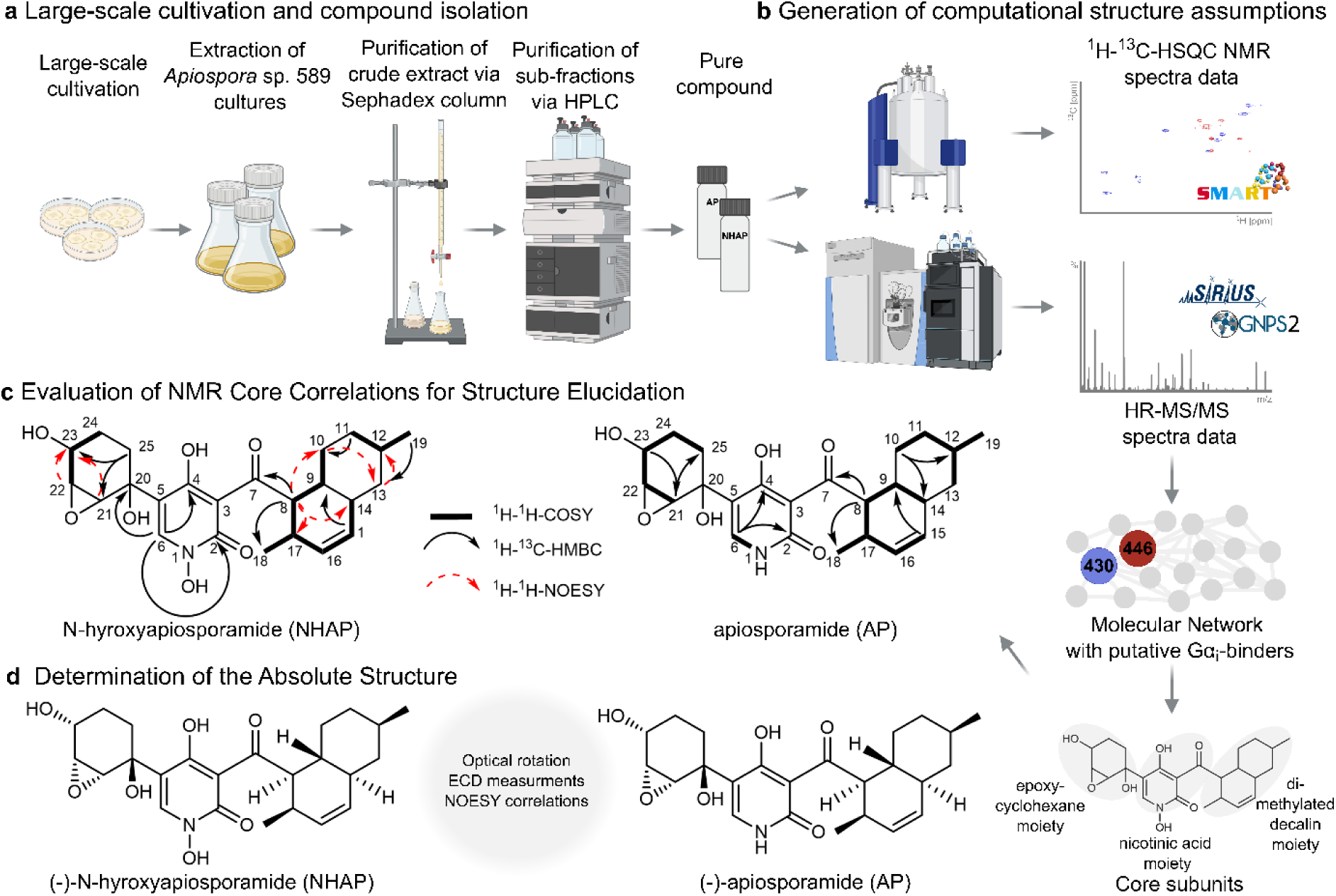
Computational-guided structure elucidation of NHAP and AP. **a** Workflow for compound extraction, isolation, and purification from *Apiospora* sp. 589. **b** Structural hypotheses generated from LC-MS/MS and ¹H-¹³C HSQC data, corroborated by molecular networking. **c** Planar structure determination via NMR analysis. **d** Stereochemical assignment using optical rotation, and ECD data.

### NHAP and AP elicit different effects on Gα_i1_

To assess functional consequences, we conducted an *in vitro* GTP-turnover assay,^58,59^ in which we assessed the ability of NHAP and AP, to inhibit G protein-catalyzed GTP depletion. (**Fig. 3a**) In this assay, NHAP and AP significantly inhibited GTP turnover of the Gα subunit Gα_i1_ **Fig. 3b**). Notably, the inhibition of GTP turnover in the presence of NHAP was observed exclusively in the presence of Gα_i1_, showing the compound’s specificity for the target protein and excluding non-specific effects on the activity of the assay enzymes. Titration of NHAP and AP resulted in a dose response curve with an IC_50_ value of approx. 43 µM and 203 µM, respectively. (**Fig. 3b**)

**Fig. 3:**
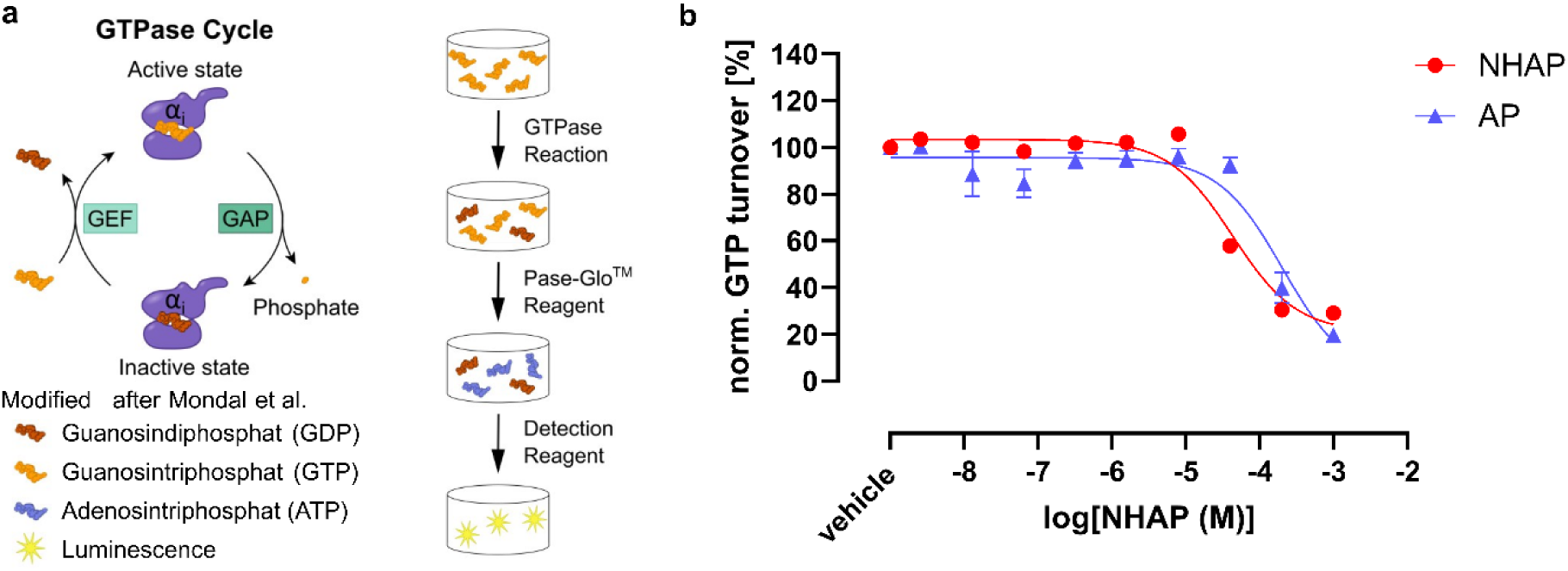
NHAP selectively inhibits GTP turnover of Gα_i1_. **a** GTPase turnover assay. **b** NHAP and AP inhibit GTP turnover of the Gα_i1_ subunit with IC_50_ values of ∼43 µM (red circles) and ∼203 µM (blue triangles), respectively. Data represent mean ± SD (n=3).

To map the potential binding site of NHAP, we employed hydrogen-deuterium exchange mass spectrometry (HDX-MS), a technique that quantifies protein conformational dynamics by measuring hydrogen-deuterium exchange rates in peptide fragments.^60^ We carried out HDXMS on the purified Gα_i1_ subunit in the presence and absence of NHAP or AP, with coverage of ∼97% of the G protein sequence. Incubation of Gα_i1_ with NHAP resulted in reduced deuterium exchange in key nucleotide-binding regions, specifically the conserved TCAT motif and the P-loop (GxGxxxG motif) after extended exchange periods (167 min) (**Fig. 4a**). In contrast, AP induced widespread HDX increases primarily in the α-helical domain, accompanied by visually observable protein precipitation, indicative of nonspecific destabilization rather than specific binding (**Fig. 4b**). Even though the changes in HDX are too limited to locate the NHAP binding site on Gα_i1_, the results are consistent with a model where NHAP engages the GDP-bound Gα_i1_ subunit in a way that leads to some stabilization of the nucleotide-binding site, while AP acts as a pan-binding destabilizer.

**Fig. 4:**
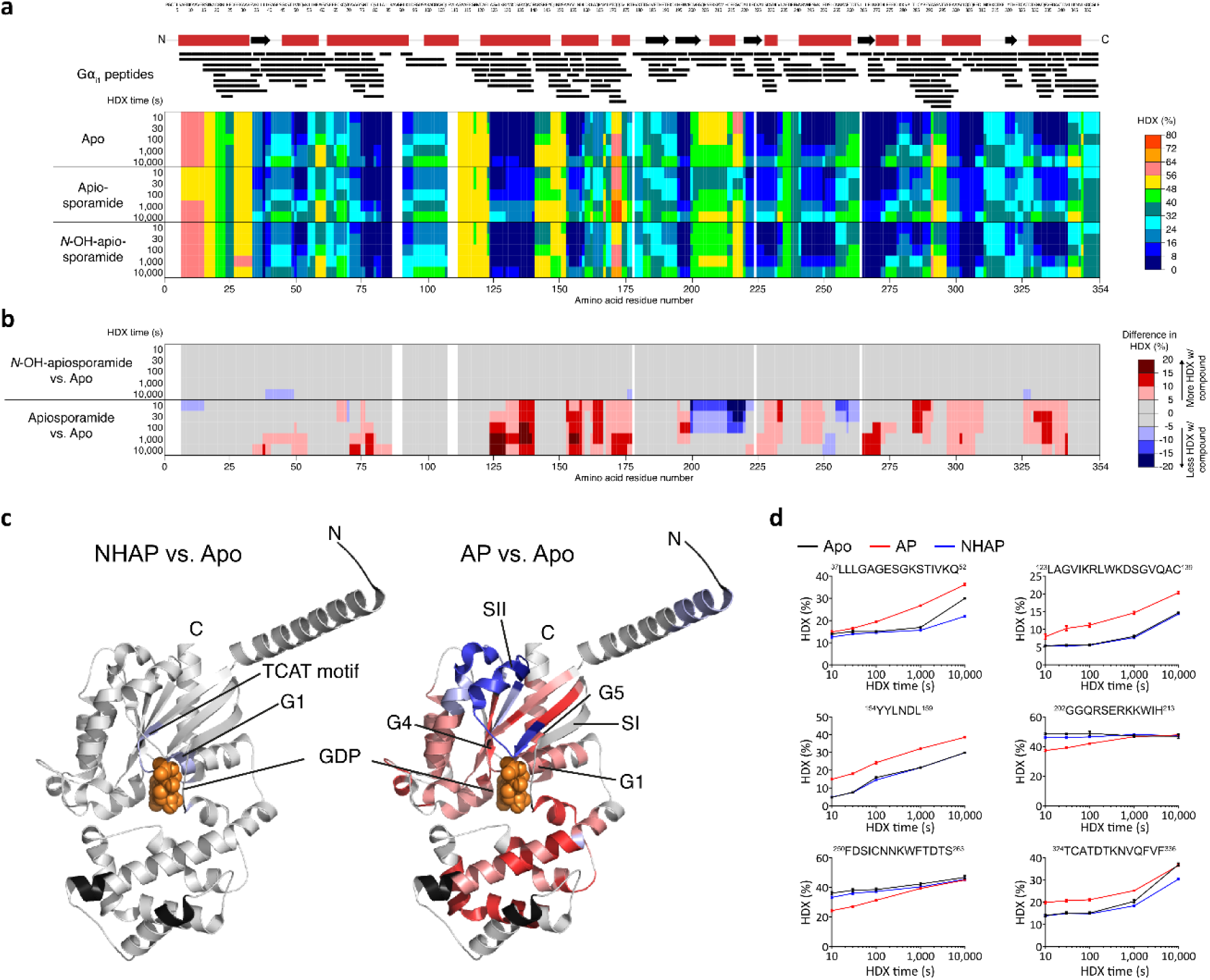
HDX-MS mapping of NHAP and AP interactions with human Gα_i1_. The secondary structure of human Gα_i1_ is schematically sketched (red boxes, α-helices; black arrows, β-strands), and black bars represent peptides derived from human Gα_i1_ identified during HDX-MS. **a** The residue-specific HDX of human Gα_i1_ either in absence (apo) or presence of compounds AP (apiosporamide) or NHAP (*N*hydroxyapiosporamide) is color-coded from 0% (blue) to 80% (red). **b** The differences in residue-specific HDX for either compound versus the apo-state are color-coded from ≤-20% (blue) to ≥20% (red) reflecting reduced and elevated HDX in the presence of either compound, respectively. **c** Projection of the HDX differences from **b** on the crystal structure of human Gα_i1_ (PDB-ID: 6CRK). The strongest change in HDX at any time-point was rendered. Black color, no data. **d** Deuterium uptake charts of representative human Gα_i1_ peptides (residue numbers in brackets). Data represent mean ± s.d. of n=3 technical replicates.

### NHAP, but not AP functionally inhibits Gi-signaling in primary cardiomyocytes

NHAP has previously been reported to display favorable bioavailability and no significant systemic toxicity in mice, whereas pronounced cytotoxic effects were observed in cancer and other proliferating cells, including HEK293 cells in our hands (**Fig. S23**).^56,61,62^ These cytotoxic effects limit the use of commonly employed proliferative cell lines for functional signaling studies, as compromised viability interferes with downstream readouts. To circumvent this limitation and assess NHAP activity in a physiologically relevant context, we turned to terminally differentiated cells.

Accordingly, we evaluated sarcomere shortening in isolated adult rat ventricular cardiomyocytes, highly specialized cells with a robust Gs- and Gi-signaling machinery regulating contractility. In this setting, Gs and Gi pathways exert well-defined and opposing roles, enabling a sensitive functional readout of Gi inhibition by NHAP. Cells were field-stimulated at 1 Hz and pre-treated with isoprenaline (Iso, 30 nM) to activate Gs-signaling, followed by addition of acetylcholine (Ach, 1 µM) to activate M2-receptor-mediated Gi-signaling (**Fig. 5a,b**). Iso induced a cAMP-mediated increase in contractility (positive-inotropic effect) which was antagonized by Ach (negative-inotropic effect) (**Fig. 5a,b**).

**Fig. 5:**
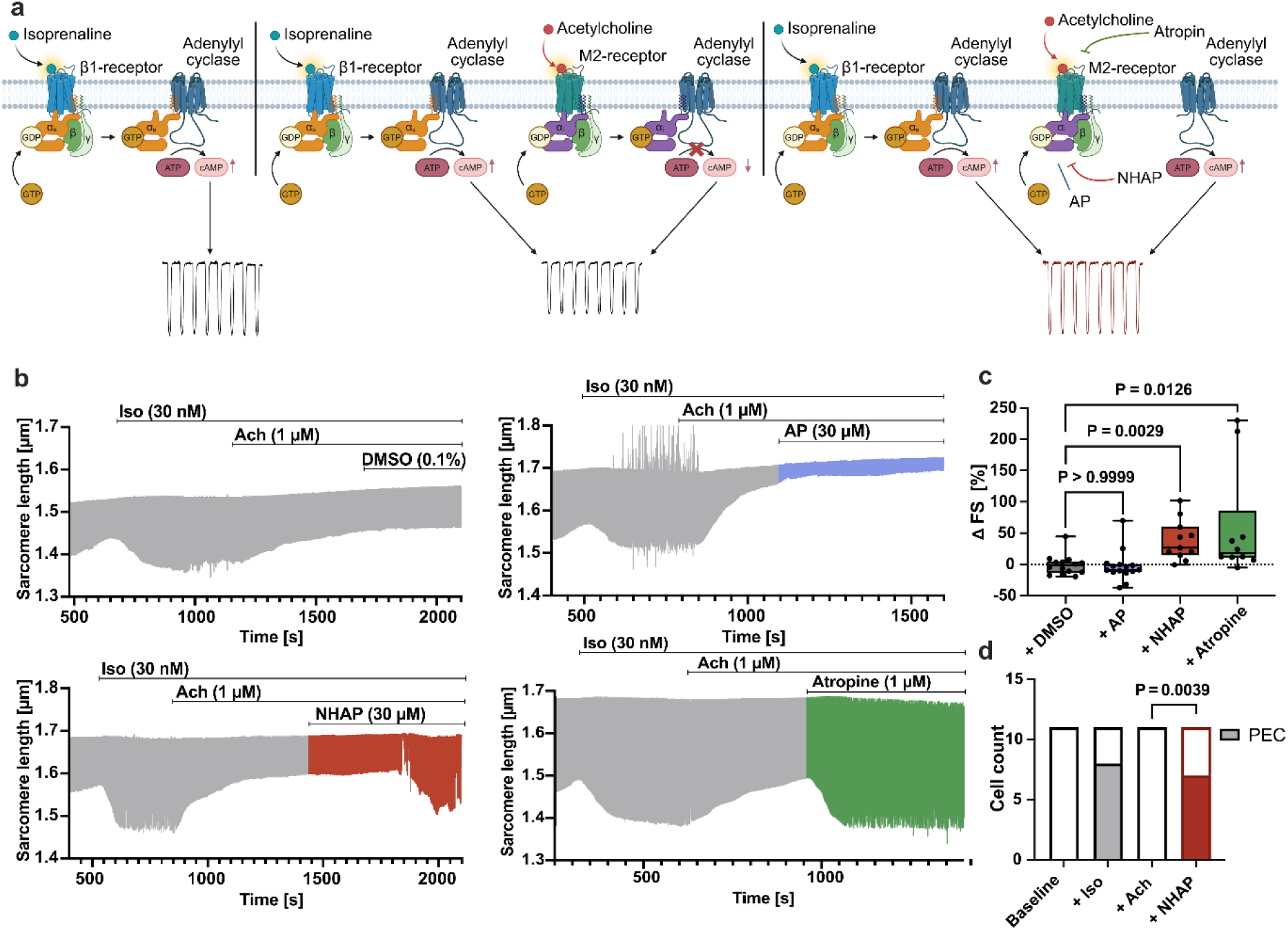
NHAP, but not AP, antagonizes acetylcholine-M2-receptor mediated effects in rat primary ventricular cardiomyocytes. **a** Scheme explaining β1and M2-receptor mediated signaling in cardiomyocytes. **b** Sarcomere shortening of rat primary ventricular cardiomyocytes after stimulation of b-adrenergic and muscarinic receptors with isoprenaline (Iso) and acetylcholine (Ach) followed by addition of DMSO, AP, NHAP or atropine, respectively. β-receptor stimulation with 30 nM Iso leads to augmented sarcomere shortening and subsequent addition of 1 µM Ach antagonizes this effect via activated M2-receptor signaling. Addition of either 0.1% DMSO or 30 µM AP has no effect on sarcomere shortening (upper left and right panel). Addition of 30 µM NHAP, on the other hand, largely reverses the Ach effect and increases sarcomere shortening presumably via direct inhibition of M2-receptor-activated Gi-signaling (lower left panel). Addition of 1 µM atropine, a muscarinic receptor antagonist, causes a similar increase in sarcomere shortening (lower right panel). **c** Change in fractional shortening (ΔFS) shows positive-inotropic effects of NHAP and atropine, whereas DMSO and AP do not alter fractional shortening significantly. Effects for DMSO, AP and atropine were calculated at 325 s after addition of the respective substance, which is the average time point of the maximal NHAP effect. (Kruskal-Wallis test). **d** Occurrence of pro-arrhythmic extra-contractions (PEC) at baseline conditions, after addition of ISO (+ Iso), subsequent addition of Ach (+ Ach) and, finally, following the application of NHAP (+ NHAP). Ach silenced the ISO-induced PEC, whereas the addition of NHAP led to the reappearance of the PEC (Fisher’s exact test). Data were obtained from the following numbers of animals (N) and cells (n) per group: DMSO: N=7, n=14; AP: N=5, n=15; NHAP: N=5, n=11; Atropine: N=5, n=10.

Addition of NHAP (30 µM) largely reversed the negative-inotropic effect of Ach (**Fig. 5b**, lower left panel). This NHAP-induced increase in contractility was observed consistently across cells (n=11), with a maximal effect reached, on average, after 325 s. Fractional shortening (FS) increased from 8.8±2.8% (pre-NHAP) to 11.5±2.5% (post-NHAP, **Fig.S24c).** In contrast, DMSO and AP (30 µM) controls did not significantly alter FS **(Fig. 5b,c).** Furthermore, NHAP restored Iso-induced pro-arrhythmic extra-contractions (PEC) that had been suppressed by Ach before (**Fig. 5d, Fig. S24b**). Iso induced PEC in most cells, likely due to calcium overload, and Ach suppressed this effect via Gi-signaling; subsequent NHAP addition re-established PEC, consistent with blockade of the anti-arrhythmic M2-Gi pathway. Similar to NHAP, the muscarinic antagonist atropine increased FS (**Fig. 5bc**) and led to re-appearance of Isoinduced PEC (**Fig.S24d**). These results demonstrate that NHAP is cell-permeable and functionally inhibits Gi-signaling in primary cardiomyocytes without acute cytotoxicity, distinguishing it from pan-G protein-binding analogs and establishing it as a suitable chemical probe for Gi pathway inhibition.

## Discussion

This study establishes native metabolomics as a powerful platform for the rapid discovery of G protein modulators from complex natural product extracts. By integrating non-targeted LCMS/MS with protein binding detection via native mass spectrometry, we identified NHAP, the first selective small molecule inhibitor of Gi-signaling, directly from complex fungal extracts. This approach bypasses the iterative cycles of traditional bioassay-guided fractionation, enabling the assignment of bioactivity to individual metabolites within hours-to-days of screening.

Our findings highlight the critical impact of subtle structural modifications on G protein selectivity. NHAP and its close analog AP differ by a single hydroxyl group yet exhibit strikingly different profiles: NHAP selectively binds to and inhibits Gα_i1_, whereas AP displays pan-Gprotein binding accompanied by protein destabilization. This mirrors the selectivity observed for the Gα_q_ inhibitor FR, where specific functional groups, especially a single side-chain hydroxyl group, dictate both binding affinity and inhibitory efficacy toward Gq-signaling.^25^ HDXMS data are consistent with modulation of the nucleotide-binding pocket, particularly the TCAT motif and P-loop region, which may underlie the observed interference with the GTPase turnover. Although additional orthogonal biophysical approaches were hampered by compound-intrinsic properties, the convergence of native MS with robust functional effects in primary cardiomyocytes provides compelling evidence for direct and functionally relevant target engagement.

The functional data in cardiomyocytes further point to a context-dependent profile of NHAP activity. NHAP has been reported to exert cytotoxic effects in cancer cells, and we likewise observed reduced viability in proliferating cell lines such as HEK293 (**Fig. S23**), which limits their utility for quantitative analysis of Gi-signaling. In contrast, terminally differentiated, postmitotic cells preserve viability under NHAP treatment and thus allow a clean functional interrogation of Gi inhibition. In adult ventricular cardiomyocytes, NHAP inhibits Gi-signaling without inducing acute cytotoxicity and even permits discrimination between G protein subtypes, revealing inhibition of Gi but not Gs pathways. These observations suggest that NHAP-mediated Gi-inhibition can be robustly assessed in physiologically relevant, nonproliferative systems, while its cytotoxic actions are likely context-dependent and at least partly independent of Gi-signaling per se. Future work will be needed to clarify the subtype specificity of NHAP within the Gα_i/o_ family members and to define the molecular determinants underlying its mechanism of action.

While NHAP represents a significant advance, future work will focus on deeper structural analyses (e.g., co-crystallography) to define the binding pose and medicinal chemistry optimization to improve potency (current IC_50_ ∼43 µM). The identification of NHAP-like analogs via genomic mining and subsequent engineering of the NHAP biosynthetic cluster offers promising avenues for deriving derivatives with enhanced properties. In conclusion, we demonstrate that native metabolomics can unlock “undruggable” targets by revealing selective modulators hidden in complex mixtures. NHAP provides the scientific community with a firstin-class pharmacological tool for dissecting Gi-signaling, paving the way for new fundamentally new therapeutic interventions against Gi-mediated diseases.

## Materials & Methods

### Marine fungi collection and taxonomy

The NHAP and AP producing marine fungus (*Apiospora* sp. strain 589) was sourced from the collection of the Institute of Pharmaceutical Biology, University of Bonn, Germany. It was isolated from a brown alga (*Fucus vesiculosus*) collected in the Bay of Lübeck during a field trip in (July 2000). A phylogenetic analysis was performed to gain a deeper understanding of the taxonomic classification of the marine fungus. The internal transcribed spacer (ITS) region was extracted using ITS primers ITS1 and ITS4. This was followed by a comparison of the identified region with reference regions in the UNITE database.^63^ The results were filtered to include hits with at least 80% coverage and at least 85% identity. The phylogram revealed a significant presence of *Apiospora* spp. in the tree beside some *Arthrinium* spp. and an *Amphichorda* sp. in closer proximity to the investigated fungus (**Figure S3**). The fungus was grown on PDA medium (24 g/ L potato/dextrose 20 g/ L agar) at 25 °C in the absence of light. Inoculation was performed with cryopreserved agar chunks from a PDA plate previously incubated with *Apiospora* sp. strain 589. The cultivation period was one month for NHAP and two months for AP. The cultures were extracted twice using an ultrasonic bath with the same volume of ethyl acetate yielding 0.16 g/L crude extract.

### Isolation and characterization of *N*-hydroxyapiosporamide and apiosporamide

NHAP and AP were initially isolated using a size exclusion column (20 × 450 mm, Sephadex LH-20, 100% methanol). The fractions were collected in 25 ml volumes. The purity of the compounds in fractions 5 and 6 was then optimized using a puriFlash 5.250P system equipped with a Phenomenex Kinetex 5 μm C18 100 Å column (21.2 × 150 mm, 35–50% acetonitrile in water over 28 min, flow rate: 22.0 mL/min, λ = 220 nm). In the final step of the procedure, the peaks eluting at minutes 15-17 (AP) and 20-22 (NHAP) were purified using an Agilent HPLC1260 series system equipped with an Eclipse XDB-C18 5µm column (9.4 x 250mm, 50- 53% acetonitrile in water over 19 min, flow rate: 3 mL/min, λ = 220 nm). Resulting AP eluting at 15 minutes and NHAP at 18 minutes with final yields of 0.25 mg/L. NMR spectra were acquired on a JEOL ECA-500 spectrometer (for NHAP) and a JEOL ECA-400S spectrometer (for AP). NMR spectra were referenced to residual solvent chloroform-*d*_1_ signals (δH 7.26 and δC 77.2). MestReNova software (version: 12.0.0-20080) was used to evaluate the NMR spectra. The UV data and CD spectra were acquired via a J-1500 CD spectrometer (Jasco) using a 1 mm path length quartz cell (Hellma Analytics). The Jasco DIP-370 was utilized in conjunction with the D-line of the sodium lamp at λ = 589.3 nm to quantify the optical rotations of the isolated compounds.

### NMR spectroscopy

***(-)N*-hydroxyapiosporamide (NHAP):** white powder; [α]^24^_D_ −75.0 (*c* 0.1, MeOH); UV (MeOH) λ_max_ (log ε) 204 (5.34), 218 (5.21), 282 (4.85), 342 (4.83) nm; ECD (0.90 mM, MeOH) λ_max_ (Δε) 204 (-5.69), 214 (-0.13), 227 (-4.73), 269 (1.64), 292 (0.95), 310 (2.06), 347 (-0.69); 1H and ^13^C NMR see **Table 1**; HRESIMS *m/z* 446.2160 [M+H]^+^ (calcd for C_24_H_32_NO_7_^+^, 446.2168)

**Table 1:**
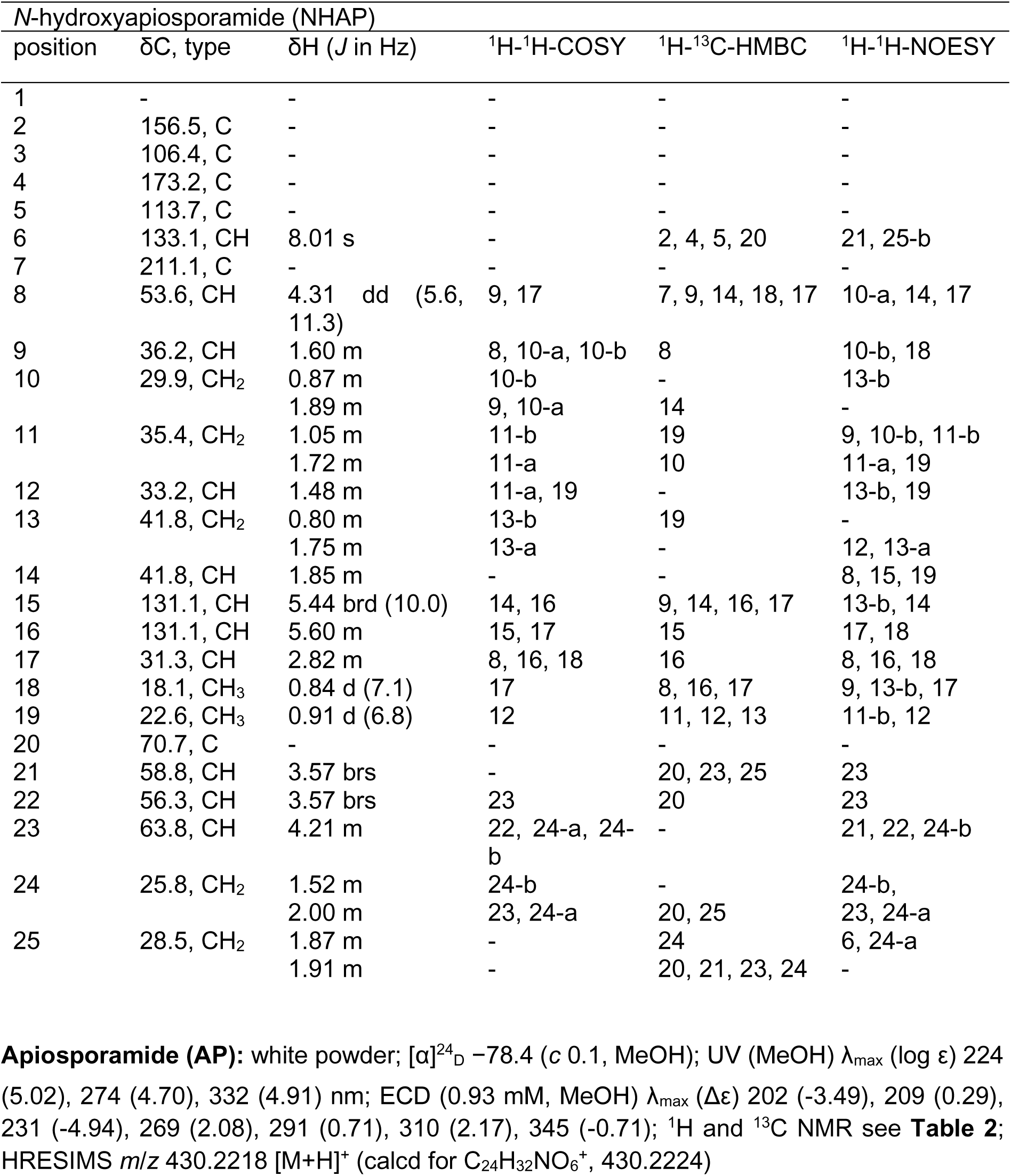
**NMR Spectroscopic Data (500/125 MHz) for *N*-hydroxyapiosporamide (NHAP) in chloroform-*d_1_***

**Table 2:** NMR Spectroscopic Data (400/100 MHz) for apiosporamide (AP) in chloroform-.

| apiosporamide (AP) |  |  |  |  |
| --- | --- | --- | --- | --- |
| position | δC, type | δH (J in Hz) | <sup>1</sup>H-<sup>1</sup>H-COSY | <sup>1</sup>H-<sup>13</sup>C-HMBC |
| 1 | - | 11.38 d (4.2) | 6 | - |
| 2 | 162.9, C | - | - | - |
| 3 | 107.9, C | - | - | - |
| 4 | 178.4, C | - | - | - |
| 5 | 116.2, C | - | - | - |
| 6 | 138.9, CH | 7.84 d (4.2) | 1 | 2, 4 |
| 7 | 211.5, C | - | - | - |
| 8 | 53.0, CH | 4.30 m | 9, 17 | 7, 17, 18 |
| 9 | 36.3, CH | 1.58 m | 8, 14 | 8 |
| 10 | 30.0, CH<sub>2</sub> | 0.86 m | 11-b | - |
|  |  | 1.87 m | 10-a | 12, 14 |
| 11 | 35.4, CH<sub>2</sub> | 1.04 m | 11-b, 12 | 19 |
|  |  | 1.72 m | 11-a, 12 | 9 |
| 12 | 33.2, CH | 1.47 m | 19 | - |
| 13 | 41.8, CH<sub>2</sub> | 0.81 m | 13-b | 19 |
|  |  | 1.72 m | 13-a, 12 | 9 |
| 14 | 42.0, CH | 1.81 m | 9 | 15 |
| 15 | 131.1, CH | 5.40 d (9.9) | 16 | 9, 14, 17 |
| 16 | 131.3, CH | 5.56 m | 15, 17 | 17 |
| 17 | 31.3, CH | 2.83 m | 18 | - |
| 18 | 18.2, CH<sub>3</sub> | 0.83 d (7.0) | 17 | 8, 16, 17 |
| 19 | 22.7, CH<sub>3</sub> | 0.90 d (6.4) | 12 | 11, 12, 13 |
| 20 | 71.0, C | - | - | - |
| 21 | 57.9, CH | 3.48 d (3.5) | 22 | 20, 25 |
| 22 | 55.4, CH | 3.57 t (3.6) | 21, 23 | - |
| 23 | 63.6, CH | 4.31 m | 24-b | 20, 21 |
| 24 | 26.1, CH<sub>2</sub> | 1.49 m | 24-b, | - |
|  |  | 2.00 m | 23, 24-a | 25 |
| 25 | 28.8, CH<sub>2</sub> | 1.81 m | 24-b | 24 |
|  |  | 1.90 m | 24-a | - |

### Plasmid DNA constructs

The human genes of Gα_i1_ and Gα_s_Δ34 (short splice variant containing residues 35-380) bearing an amino terminal 8xHis-tag and a HRV 3C cleavage site were subcloned into the plasmids pET21a and pET28a (Merck), respectively, for bacterial expression. For the expression of the heterotrimeric Gi1 in insect cells, the human Gα_i1_ subunit was cloned into pVL1392 (Expression Systems), and the human Gβ1 and Gγ2 subunits were both cloned into a single modified pVL1392 dual vector as described previously.^64^ A 6xHis-tag and a HRV 3C protease site were attached to the N-terminus of Gβ1 for purification. To generate baculoviruses for the expression of non-lipidated Gβ1γ2 heterodimer, the carboxy terminal cysteine (C68) of the Gγ2 subunit was mutated to serine in the modified pVL1392 dual vector using Quick-change mutagenesis.

### Expression and purification of Gα subunits

Gα subunits (human Gα_i1_ and Gα_s_Δ34) were expressed and purified as previously described.^65^ Briefly, the protein was expressed in Rosetta 2 (DE3) cells (EMD Millipore) using pET21a (Gα_i1_) or pET28a (Gα_s_Δ34). Cells were grown in Terrific Broth to OD600 of 0.6, and protein expression was induced by addition of 0.5 mM IPTG. After 15 h of incubation at 20 °C, cells were harvested and resuspended in lysis buffer (50 mM HEPES pH 7.5, 100 mM sodium chloride, 1 mM magnesium chloride, 50 μM GDP, 5 mM β-mercaptoethanol, 5 mM imidazole, and protease inhibitors). Cells were disrupted by sonification using a 50% duty cycle, 70% power for four times 60 s on ice. Intact cells and cell debris were subsequently removed by centrifugation and the supernatant was incubated with Ni-NTA resin for 1.5 h at 4 °C. The Ni-NTA resin was washed multiple times in batch with lysis supplemented with 20 mM imidazole and then loaded into a wide-bore glass column. After washing the resin with 10 CV with the same buffer, containing 20 mM imidazole, the protein was eluted with lysis buffer supplemented with 200 mM imidazole. The eluted protein was dialyzed overnight in dialysis buffer (20 mM HEPES pH 7.5, 100 mM sodium chloride, 1 mM magnesium chloride, 10 μM (Gα_i1_) or 20 µM (Gα_s_Δ34) GDP, 5 mM β-mercaptoethanol, and 5 mM imidazole). The amino terminal His-tag was cleaved by adding 1:500 w/w HRV 3C protease into the dialysis bag. Uncleaved protein, cleaved His-tag, and HRV 3C protease were subsequently removed by loading the protein on Ni-NTA resin and collecting the flow-through. The protein was concentrated and run on a Superdex 200 10/300 GL column in size exclusion buffer (20 mM HEPES pH 7.5, 100 mM sodium chloride, 1 mM magnesium chloride, 10 μM (Gα_i1_) or 20 µM (Gα_s_Δ34) GDP, and 100 uM TCEP). The monomeric peak fractions were pooled, concentrated and frozen in N_2_ after the addition of 20% (v/v) glycerol. The protein was stored at -80°C until further use. Purification of G_11_ was performed as previously described by Mühle et al.^29^

### Micro-flow LC-MS/MS data acquisition – Metabolomics

Analyses were performed using a Vanquish UHPLC system (Thermo Fisher Scientific, Bremen, Germany) coupled to a Q-Exactive HF quadrupole orbitrap mass spectrometer (Thermo Fisher Scientific, Bremen, Germany). A sample volume of 2 µL was injected onto a C18 reversed-phase core-shell micro-flow column (Kinetex C18, 150 x 1 mm, 1.8 um particle size, Phenomenex, Torrance, USA). Solvent A (H_2_O + 0.1% formic acid (FA)) and solvent B (acetonitrile (ACN) + 0.1% FA) were used as the mobile phase. The flow rate was set to 60 µL/min. The gradient was applied: 10–90% B between 0 and 5 min, followed by 90–99% B between 5 and 10 min, followed by a 3-min washout phase at 99% B and a 5-min re-equilibration phase at 10% B.

MS/MS spectra were recorded in positive ion mode using data-dependent acquisition (DDA). Electrospray ionization (ESI) settings were as follows: capillary temperature 253 °C, sheath gas flow 46.25 auxiliary gas flow 10.63 sweep gas flow rate 2.13 and spray voltage to 3.5 kV. The inlet capillary was heated to 406 °C and the S-lens level was set to 50 V applied. MS scan range was set to 200 - 2000 m/z with a resolution of 120,000 with one micro-scan. With an automatic gain control (AGC) offset to 1e^6^, the maximum ion injection time was 100 ms. For MS/MS spectra, one micro-scan was recorded per duty cycle at a resolution of 15,000. For MS/MS scans the AGC target was 5e^5^ ions with a maximum ion injection time of 50 ms. The isolation window for the precursor-ion was set to *m/z* 1. We applied a stepped normalized collision energy from 25 to 35 to 45. 10 s was set for the dynamic precursor exclusion.

### Native metabolomics

Gα_i1_ protein was rebuffered to 100 mM ammonium acetate (pH 7.5). By using Amicon® Ultra Centrifugal Filters, 4 cycles of ultrafiltration at 4 °C. The final concentration of the Gα_i1_ protein was set to 0.9 mg/mL. To generate native like conditions for the protein, a make-up flow via the quaternary pump was constantly infused with a flow rate of 200 µL/min at 50% of 100 mM ammonium acetate. Due to a syringe pump the protein-solution was constantly infused with a flow rate of 2 µL/min. A sample volume of 2 µL was injected and an identical metabolomics run was conducted for the chromatographic separation of the crude extract, but without infusion of protein-solution.

At a resolution of 15,000 (m/z 200) the MS scan range was set to 2,500 to 5,000 m/z. All-Ion Fragmentation (AIF) was performed with a collision energy (CE) of 10 eV. AGC target was set to 3e6 and the maximum ion injection time was set to 2000 ms. Spray voltage was applied to 3 kV with S-lens RF level at 70 and heated capillary temperature at 253 °C. The sheath gas flow rate was set to 40, the aux gas flow rate to 10 and the sweep gas flow rate to 2. The probe heater temperature was set to 150 °C.

### Native mass spectrometry

#### Selectivity test with NHAP/AP

Purified Gα_i1_, Gα_s_, or G11-heterotrimer were buffer exchanged in 100 mM ammonium acetate prior to native MS measurements. The compounds NHAP and AP were analyzed at a concentration ten times higher than that of the respective G-protein.

All native MS analyses were carried out using a spray voltage of 3 kV with S-lens RF level at 70. The probe heater temperature was set to 50 °C and heated capillary temperature to 253 °C. The sheath gas flow rate was set to 20, the aux gas flow rate to 7 and the sweep gas flow rate to 2. A higher collision energy of 10 was conducted, respectively.

Denaturation measurement: A solution of 40 µM Gα_i1_ was incubated with an equal volume of 7 M urea.^66^ The urea was then exchanged using 100 mM ammonium acetate. After buffer exchange an excess of 10-fold NHAP/AP was added prior to the native MS measurement.

#### Native metabolomics data analysis

Data were analyzed with Xcalibur 2.2. for deconvolution of the data, quantification and determination of the charge states of Gα_i1_, Gα_s_, G_11_-heterotrimer and FR-sensitive Gα_i1_ Unidec^67^ software was used.

### Calculation of IC_50_ values

The IC_50_ value for NHAP in the GTP turnover assay was determined by using a three-parameter sigmoidal curve-fit in Prism 10 (GraphPad), based on the equation Y=Bottom + (Top-Bottom)/(1+10((LogIC50-X)*HillSlope)).

### Hydrogen/deuterium exchange mass spectrometry (HDX-MS)

Stand-alone human Gα_i1_ was isolated as described above and supplemented with 2.5% (v/v) of 100 mM NHAP, AP or DMSO prior HDX-MS.

The preparation of individual HDX reactions was assisted by a two-arm robotic autosampler (LEAP Technologies) as described previously.^68^ In brief, HDX reactions were initiated by 10- fold dilution of human Gα_i1_ (20 µM) with buffer (20 mM Tris-Cl pH 8.0, 2 mM MgCl_2_, 20 µM GDP, 2.5% (v/v) DMSO) prepared in D_2_O. After incubation for 10, 30, 100, 1,000 or 10,000 s at 25 °C, the HDX reaction was halted through mixing with an equal volume of quench buffer (400 mM KH_2_PO_4_/H_3_PO_4_, 2 M guanidine-HCl; pH 2.2) temperated at 1 °C. Non-deuterated samples were generated through 10-fold dilution with buffer prepared with H_2_O and treated similar. Immediately afterwards, 100 µL of the resulting mixture were injected into an ACQUITY UPLC M-Class System with HDX Technology.^69^ The sample injections were washed out of the injection loop (50 μL) with H_2_O + 0.1% (v/v) formic acid at a flow rate of 100 μL/min, and proteins digested by passing them through a cartridge (2 mm x 2 cm, 12 °C) filled with bead-immobilized protease, i.e., porcine pepsin or a 1:1 mixture of protease type XVIII from *Rhizopus* sp. and protease type XIII from *Aspergillus saitoi*. Resulting peptides were collected on an AQUITY UPLC BEH C18 VanGuard column (2.1 x 5 mm, 1.7 µm; Waters) at 0.5 °C. After 3 min of digestion, the peptide trap was placed in line with an ACQUITY UPLC BEH C18 column (1.0 x 100 mm, 1.7 µm; Waters), and peptides eluted at 0.5 °C with a gradient of H_2_O + 0.1% (v/v) FA (solvent A) and ACN + 0.1% (v/v) FA (solvent B) at 30 µL/min as follows: 0-7 min: 95-65% A; 7-8 min: 65-15% A; 8-10 min: 15% A; 10-11 min: 5% A; 11-16 min: 95% A. Peptides were ionized by electrospray ionization (250 °C capillary temperature, 3.0 kV spray voltage) and mass spectra acquired on a G2-Si HDMS with ion-mobility separation (Waters) in positive-ion mode from 50 to 2,000 m/z using enhanced high-definition MS (HDMS^E^) and high-definition MS (HDMS) modes for non-deuterated and deuterated samples, respectively.^70,71^ Lock-mass correction was performed with [Glu1]-fibrinopeptide B standard (Waters). During peptide separation on the C18 column, the protease column was washed 3 times with 80 µL each of 0.5 M guanidine-HCl in 4% (v/v) CAN and blank injections performed between each sample to reduce peptide carry-over. For each protein state and time-point, three individual HDX reactions were measured per protease.

Identification of peptides and analysis of their deuterium incorporation were conducted with ProteinLynx Global SERVER v. 3.0.1 (PLGS; Waters) and DynamX v. 3.0 (Waters), as described previously.^68^ Data obtained from digestion with porcine pepsin or the fungal protease proteases were combined to improve peptide coverage and redundancy. Only peptides with an intensity >10,000 counts, 5-40 residues, *≥* 2 products with *≥* 0.05 products/residue, *≤* 25 ppm mass error and retention time tolerance of 0.5 minutes were analysed. All spectra were manually inspected and, if necessary, individual samples or/and peptides excluded from the analysis.

For rendering the residue-specific HDX differences from overlapping peptides (see Supplementary Dataset 1), the shortest peptide covering a certain residue was used; where multiple peptides were of the shortest length, the peptide with the residue closest to the peptide’s C-terminus was utilized.

### GTP turnover assay

GTP turnover of Gα_i1_ was measured using the GTPase-Glo™ assay (Promega) as described previously^58^ with the following modifications: For the initial activity screening of G protein binders on the GTP turnover of Gα_i1_, the reaction buffer consisted of 20 mM HEPES pH 7.5, 100 mM sodium chloride, 10 mM magnesium chloride, and 5 µM GTP. Before starting the reaction, 2 µM Gα_i1_ was incubated in the presence of 2 µM or 200 µM hit compound or equal volumes of the corresponding solvent (buffer or methanol) for 30 minutes on ice in a buffer containing 20 mM HEPES pH 7.5, 100 mM sodium chloride, and 20 mM magnesium chloride. To test for unspecific activity of the compounds on the assay enzymes, the GTP turnover assay was also performed in the absence of Gα_i1_. The reaction was initiated by the addition of an equal volume of buffer (20 mM HEPES pH 7.5, 100 mM sodium chloride) containing 10 µM GTP to reach final concentrations of 1 µM Gα_i1_, 10 mM magnesium chloride, 1 or 100 µM hit compound, and 5 µM GTP. After 1 hour incubation at room temperature (RT), reconstituted GTPase-Glo reagent was added to the samples and incubated for an additional 30 minutes at RT. Luminescence was measured after the addition of detection reagent and subsequent incubation for 10 minutes at room temperature (RT) using a TECAN Spark Multimode Microplate reader, and the data were plotted using GraphPad Prism. For collecting dose-response curves of NHAP, the reaction buffer was changed to 20 mM Tris/HCl pH 8.0, 100 mM sodium chloride, 10 mM magnesium chloride, and 5 µM GTP to increase the solubility of

NHAP. Before starting the reaction by the addition of GTP, as described above, 2 µM Gα subunit (Gα_i1_) or G_i1_ heterotrimer (Gα_i1_Gβ_1_Gγ_2_) was incubated with different concentrations of NHAP (25.6 nM, 128 nM, 640 nM, 3.2 µM, 16 µM, 80 µM, 400 µM, and 2 mM) or solvent (DMSO) for 30 minutes on ice. After 2 h incubation of the assay at RT, reconstituted GTPase-Glo reagent was added to the samples and incubated for 30 minutes at RT. Measurement of the luminescence and data analysis were performed as described above.

### Cell Viability Assay

Cell viability was assessed using the CellTiter Blue® Cell Viability Assay Kit (Promega®). A total of 6,250 HEK293 were seeded into 384 well black clear-bottom plates (BD Falcon®). 12 h after seeding the indicated concentrations of NHAP/AP diluted in culture medium, medium alone, or DMSO-containing medium were added to the cells. 24 h after seeding 6 µl of CellTiter-Blue® Reagent were added to the cells and incubated for 2 h at 37 °C. Fluorescence intensity was measured at 590 nm (excitation: 560 nm, cutoff: 570 nm) using a FLEXStation 3 (Molecular Devices). Baseline fluorescence of the CellTiter Blue dye was subtracted, and data were normalized to fluorescence values obtained from cells incubated with medium only. Data are shown as mean ± SEM of at least 3 independent experiments performed in triplicate. Statistical analysis was conducted using 2-way ANOVA with multiple comparisons in Graph Pad Prism.

### Dynamic Mass Redistribution (DMR)

Label-free whole-cell biosensing based on detection of dynamic mass redistribution (DMR) was performed as previously described in detail.^72,73^ Briefly, HEK 293 cells were seeded into fibronectin-coated 384-well Epic biosensor plates (Corning) at a density of 18,000 cells/well and cultured under standard conditions. 12 h after seeding, NHAP was added to final concentrations of 100 µM or 300 µM per well, or cells were treated with DMSO-containing medium as control. After an additional 12 h of incubation, cells were washed twice with HBSS supplemented with 20 mM HEPES and equilibrated for 60 min in the Epic reader (Corning). After equilibration, measurements were initiated, and Carbachol or EGF were added to final concentrations of 100 µM and 1 µM, respectively, using a semi-automated liquid handling system (Cybio-Selma, Analytik Jena). DMR-responses triggered by the compounds were recorded for at least 60 min at 37 °C. Raw data were buffer-corrected by subtracting the mean signal obtained after addition of HBSS+HEPES and normalized to the maximal response induced by 100 µM carbachol. Experiments were performed in triplicate, and data are presented as mean ± SEM of at least 2 independent experiments. Statistical analysis was performed on the maximal responses of the means of each individual experiment using 2-way ANOVA with multiple comparisons in Graph Pad Prism.

### Cardiomyocytes

Ventricular cardiomyocytes from hearts of Wistar-Kyoto rats (WKY) at the age of 12-20 weeks were isolated enzymatically by means of the Langendorff perfusion technique as described previously.^74^ Isolated cardiomyocytes were seeded on glass plates and mounted into a recording chamber on the stage of an inverted microscope. Myocytes were superfused with normal Tyrode’s (NT) solution containing (mM): 140 NaCl, 5.4 KCl, 1.5 CaCl_2_, 0.5 MgCl_2_, 10 HEPES, 10 glucose (pH 7.4). Myocytes were electrically-stimulated via two parallel platinum wires at 1 Hz. Only cells with clear cross striations and rhythmic sarcomere shortenings (under baseline conditions) were chosen for experiments. Moreover, cells had to respond to both isoprenaline and acetylcholine with clear increases and decreases, respectively, in sarcomere shortenings to be included in the analysis (see below). All experiments were conducted at room temperature.

Unloaded sarcomere shortening was recorded from a rectangular region-of-interest covering part of the myocyte using an IonOptix setup (IonOptix Ltd., Dublin, Ireland). Sarcomere shortening was analyzed using the IonWizard Software (Version 6.6). Fractional shortening (FS), i.e. the shortening amplitude normalized to diastolic sarcomere length, is provided as a measure of the magnitude of shortening or the strength of contraction.

Following an initial period of several minutes during which cells were superfused with NT solution (and exhibited some rundown in contractility), solutions containing isoprenaline (Iso), Iso plus acetylcholine (Ach), and, finally, Iso plus Ach plus either DMSO, AP, NHAP or atropine were applied. Solutions were supplied from reservoirs connected to a central plastic tube which led into the recording chamber. Solution change was controlled electronically. The lag time, i.e. the time period from the switch of the solution until the new solution reached the cell in the recording chamber, was approximately 150-180 s. Exposure of the cells to the pharmacological substances applied is indicated in the figures after a lag time of 150 s.

Aqueous stock solutions of Iso (1 mM with 5 mM ascorbic acid), Ach (100 mM) and atropine (10 mM) were prepared freshly on the day of the experiments. Stock solutions of AP (30 mM), NHAP (30 mM) were prepared in DMSO and stored at -20 °C.

## Data availability

The NMR data for AP and NHAP have been deposited in the Natural Products Magnetic Resonance Database (NP-MRD). All raw (.raw) and centroided (.mzML) mass spectrometry data and processed data feature tables (.csv) are publicly available via the MassIVE repository (massive.ucsd.edu) identifier MSV000102161.

HDX-MS data are provided in the spreadsheet Supplementary Dataset 1, and the full HDX-MS raw dataset been deposited to the ProteomeXchange Consortium via the PRIDE partner^75^ repository with the dataset identifier PXD080443.

Note for the editor/reviewers: MS, NMR, and HDX-MS data will be submitted to the above-mentioned FAIR-compliant databases upon acceptance of the manuscript.

## Supporting information

SupplementaryDataset_HDX

## Acknowledgments

For technical support with NMR measurements, we thank Stefan Newel. We acknowledge support by the German Research Foundation (DFG) through the core facility for HDX-MS (projects 260989694 and 324652314 to G.B.), Q-exactive LC-MS/MS measurements (INST 160/763-1 FUGG to U.L.) and project 290827466/FOR2372 (grants 216619161 to G.S., and 418513893 to X.D.). R.R. was supported by Germany’s Joint Federal and State Program Supporting Early-Career Researchers (WISNA) and by a material cost grant by Fonds der Chemischen Industrie. E. K. was funded by the Deutsche Forschungsgemeinschaft (DFG, German Research Foundation) - Project-ID 494832089 - RTG; L.J. is a member of RTG 2873 and was funded through this program.

The authors declare no competing financial interest.

