## Supporting Information for "Discovery of a Cell-Permeable Gi-Signaling Inhibitor"

*These authors contributed equally.

### Table of Figures

### List of Table

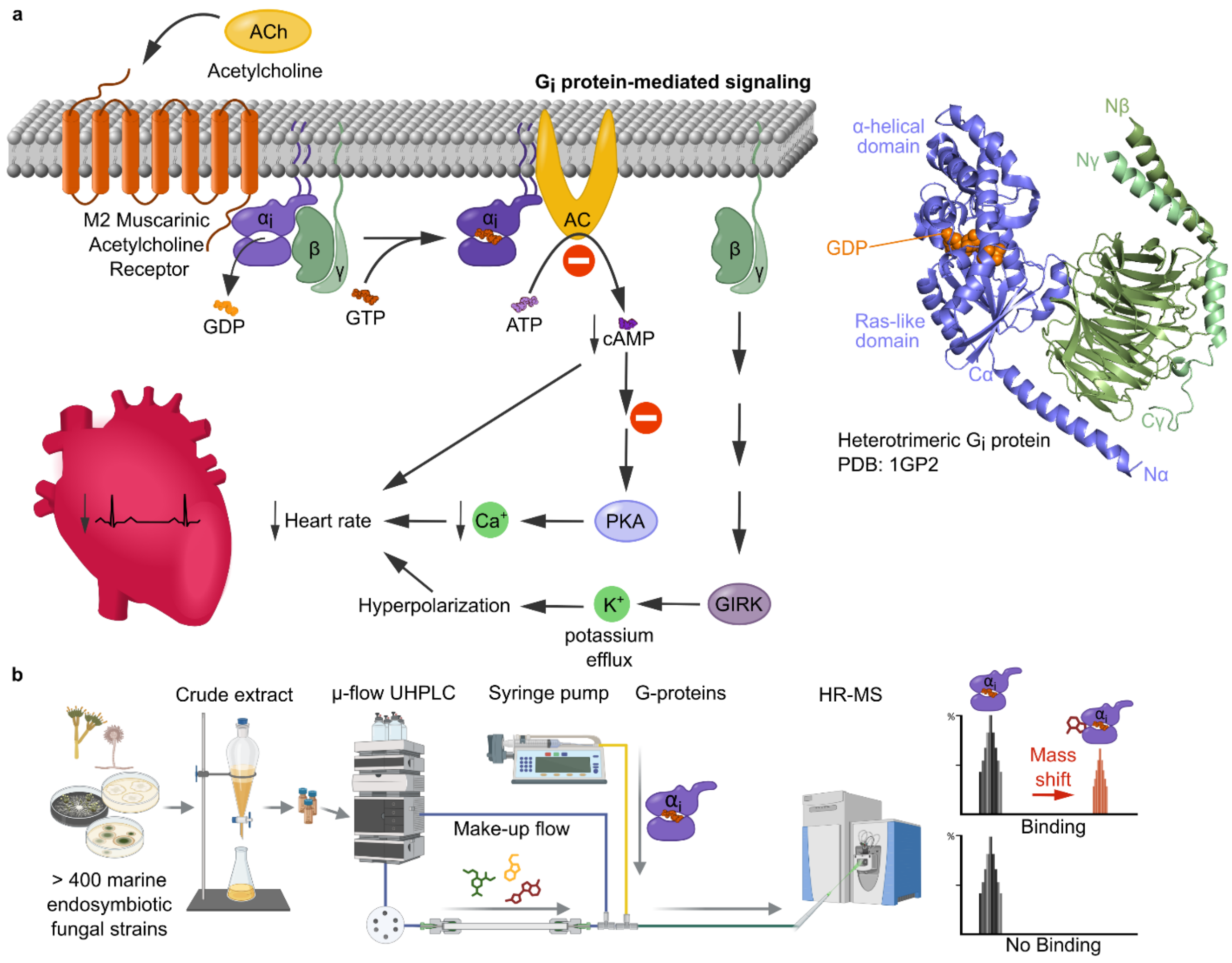

**Supplementary Figure 1: Gi protein mediated signaling and optimization of native metabolomics for the discovery of $G\alpha_{i1}$ protein binders. ........................................................**
**a** Binding of the ligand acetylcholine to the M2 muscarinic receptor triggers a complex signaling cascade in the cell that induces a conformational change in the receptor, which in turn leads to activation of the heterotrimeric G protein. The Gi heterotrimer consists of the subunits $G\alpha_i$, Gβ and Gγ. By binding Guanosine triphosphate (GTP) instead of Guanosine diphosphate (GDP) to $G\alpha_i$, $G\alpha_i$ dissociates from Gβγ. Activation of $G\alpha_i$ leads to inhibition of the cyclic adenosine monophosphate (cAMP) signaling pathway via adenyl cyclase (AC). This activates protein kinase A (PKA), which phosphorylates a variety of target proteins and thereby modulates signal transduction pathways associated with cell growth, cell differentiation and other cellular processes. The structure of the heterotrimeric Gi protein was created based on the coordinates from the PDB file 1GP2 and visualized using pymol. **b** Native Metabolomics setup: A crude extract is separated by microflow ultra-high performance liquid chromatography (µ-flow UHPLC). After chromatography, the pH of the eluent is adjusted with ammonium acetate via the make-up pump to conditions similar to the physiological environment. At the same time, the protein of interest is continuously injected orthogonally. The protein binding complexes are then measured by mass spectrometry (MS). In parallel, a metabolomics run is measured in which high-resolution UHPLC-MS/MS acquisition is performed without protein infusion. Optimization: The acetonitrile concentration has been reduced in order to denature as little protein as possible and still obtain good chromatography. By testing different G-protein concentrations, we found the optimal signal-to-noise ratio. We were able to reduce the consumption from 100 µM G-protein to 20 µM G-protein.

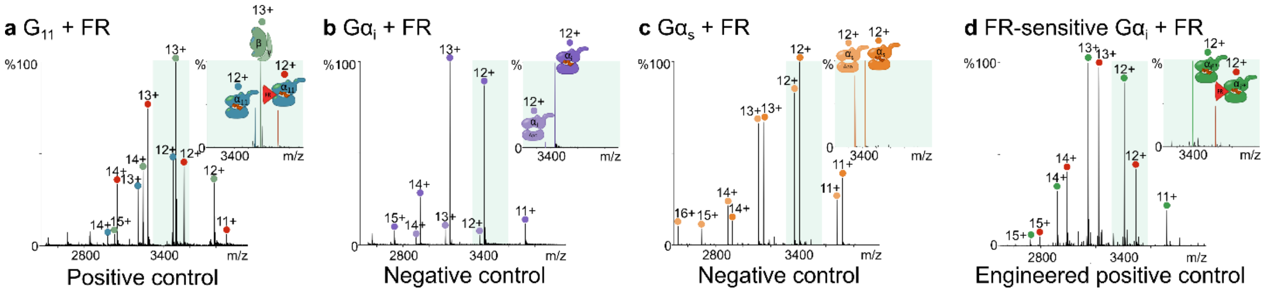

**Supplementary Figure 2: Native mass spectrometry experiments as proof-of-concept for different G-protein families.**

**a** positive control: Native mass spectra of G11 and FR900359 (FR, a highly selective and potent G$\alpha_{q/11/14}$ inhibitor). **b** negative control: Native mass spectra of G$\alpha_{i1}$ and FR. **c** negative control: Native mass spectra of G$\alpha_s$ and FR. **d** engineered positive control: Native mass spectra of FR-sensitive G$\alpha_{i1}$ and FR.

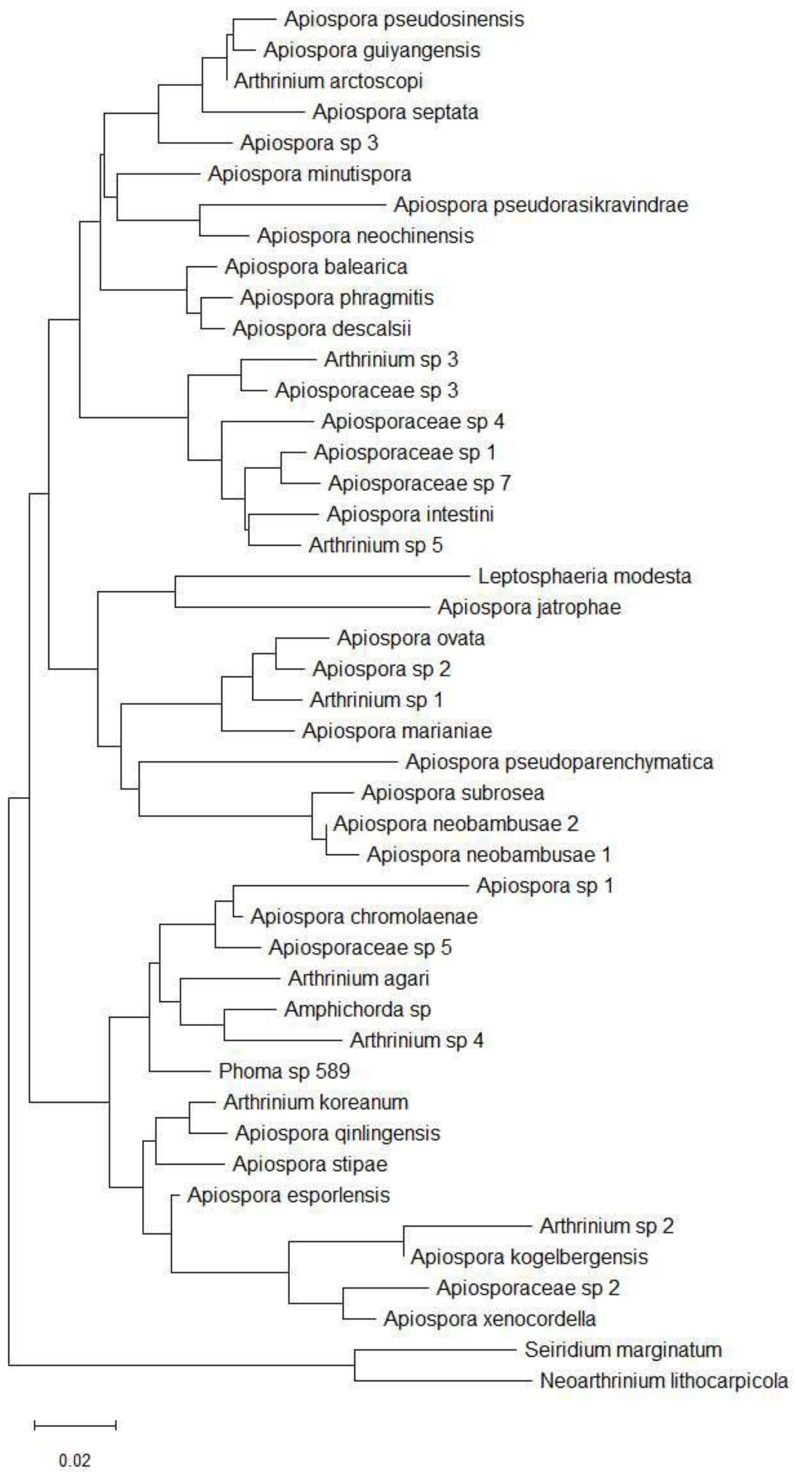

**Supplementary Figure 3: Phylogenetic tree of ITS-regions with at least 80% coverage and at least 85% identity in comparison with ITS-region of *Apiospora* sp. 589 (formerly known as *Phoma* sp. 589) found in the UNITE database.[1]**

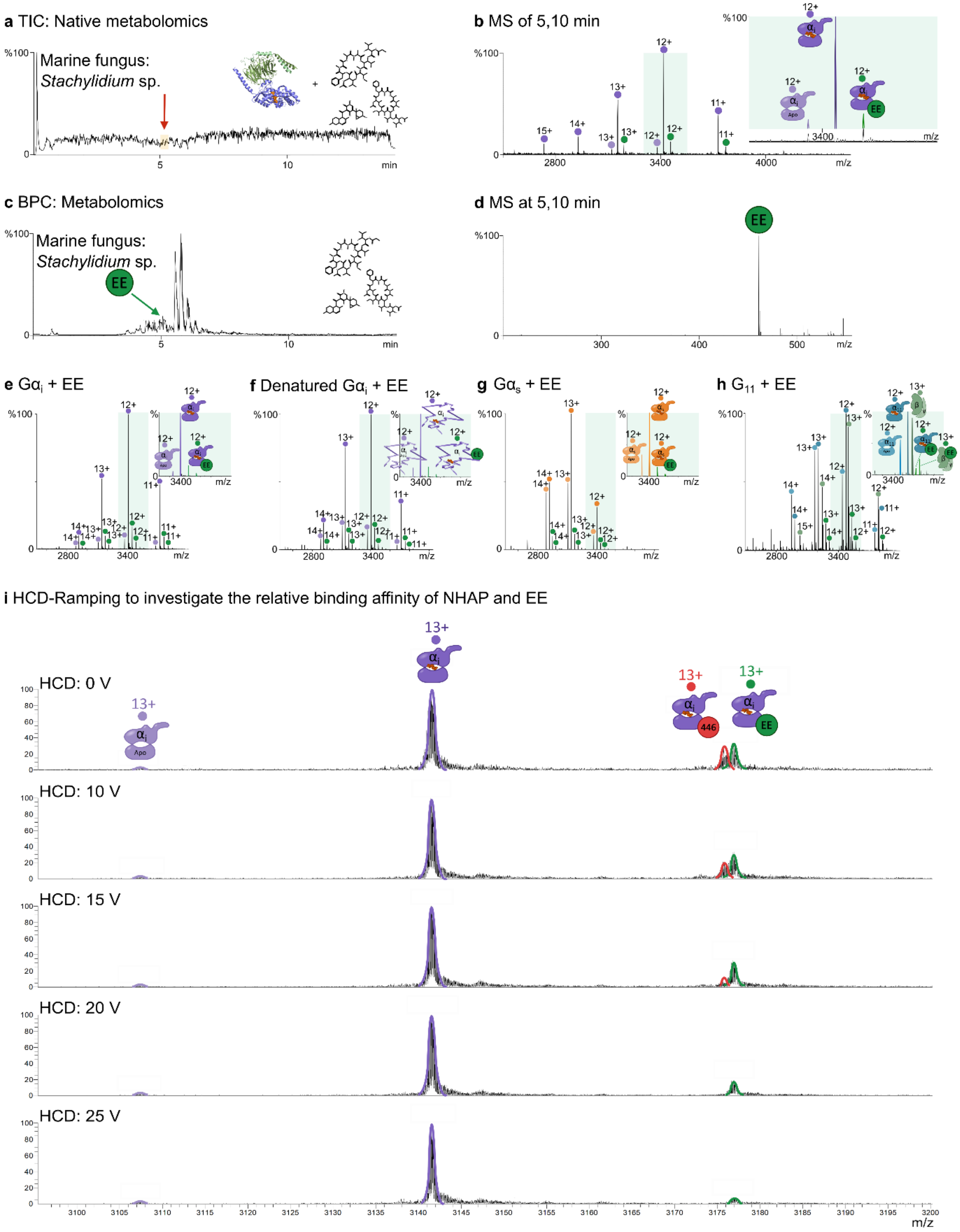

**Supplementary Figure 4: Discovery of Gα$_i$ binder from *Stachylidium bicolor* crude extract.** **a** Total ion chromatogram (TIC) of the native metabolomics run with Gα$_{i1}$ and the crude extract of *Stachylidium bicolor* 293 K04. The total ion chromatogram shows a drop at a retention time of 5.10 min. **b** Native mass spectrum of Gα$_{i1}$ at the retention time of 5.10 min. The signal of the Apo-Gα$_i$

protein is shown in light purple (charge state 12+ is shown) and the signal of the GDP-G$\alpha_{i1}$ protein in dark purple (charge states 11+ to 15+ are shown). A clear mass shift (green signal, charge states 11+ to 13+) can be seen with the signal of the GDP-G$\alpha_{i1}$ protein binding to endolide E (EE). **c** Base peak chromatogram (BPC) of the metabolomics run of the crude extract of *Stachylidium bicolor* 293 K04. At the same retention time of 5.10 min, a dominant peak with a mass of *m/z* 461 can be recognized. **d** MS1 spectrum at retention time of 5.10 min (*m/z* 461). **e** Native mass spectra of G$\alpha_{i1}$ and (EE, dark green) (10x excess). Mass shift of GDP-G$\alpha_{i1}$-EE in green. **f** Native mass spectra of denatured G$\alpha_{i1}$ and GDP-G$\alpha_{i1}$ (charge states in 12+). Binding of EE to GDP-G$\alpha_{i1}$ (in green). **g** Native mass spectra of G$\alpha_s$ and EE (10x excess). Mass shift in green **h** Native mass spectra of $G_{11}$ (charge states of GDP-G$\alpha_{11}$ in light blue and Apo-G$\alpha_{11}$ in blue in 11+ to 15+ and Gβγ in 12+ to 15+ in green) and EE (5x excess, dark green, charge states of Gβγ-EE in 12+ to 14+). Mass shift in green to the α-subunit and dark green to the βγ-subunit. **i** We used HCD-ramping to investigate the relative binding affinity of NHAP and EE (as a negative control) to G$\alpha_{i1}$. We used HCD-ramping specifically to dislodge the molecules from the protein, allowing us to observe the dissociation process of the protein-binder complexes in detail. By gradually increasing the collision energy (0 - 25 V), we gained valuable insight into the binding behavior of NHAP and EE in the presence of G$\alpha_{i1}$ and were able to better understand their potential as binders.

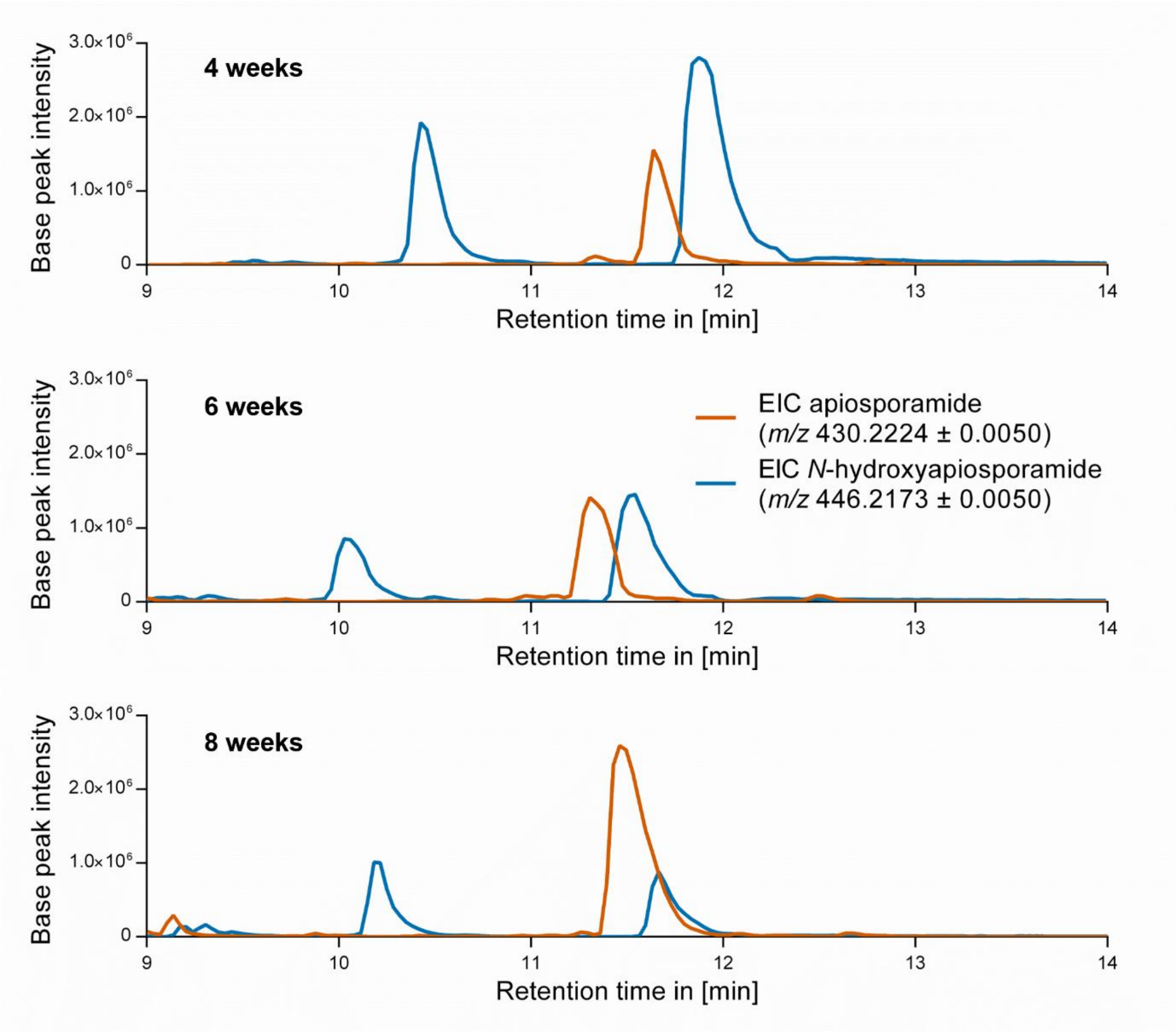

**Supplementary Figure 5: Time course experiments: extracted ion chromatograms of AP (orange) and NHAP (blue) at different extraction times.**

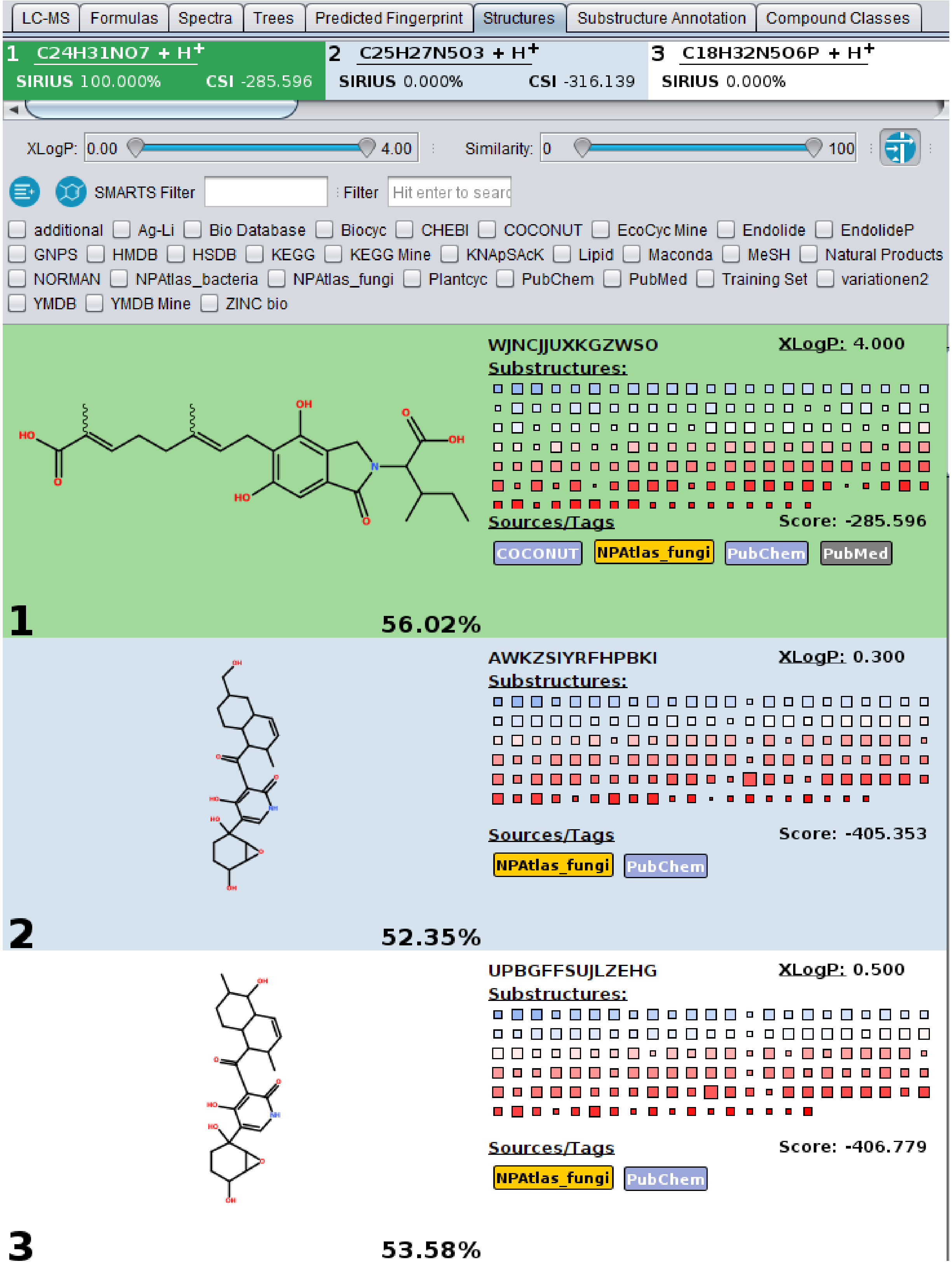

**Supplementary Figure 6: SIRIUS[2] molecular formula and CSI:FingerID[3] structure assumptions for NHAP.**

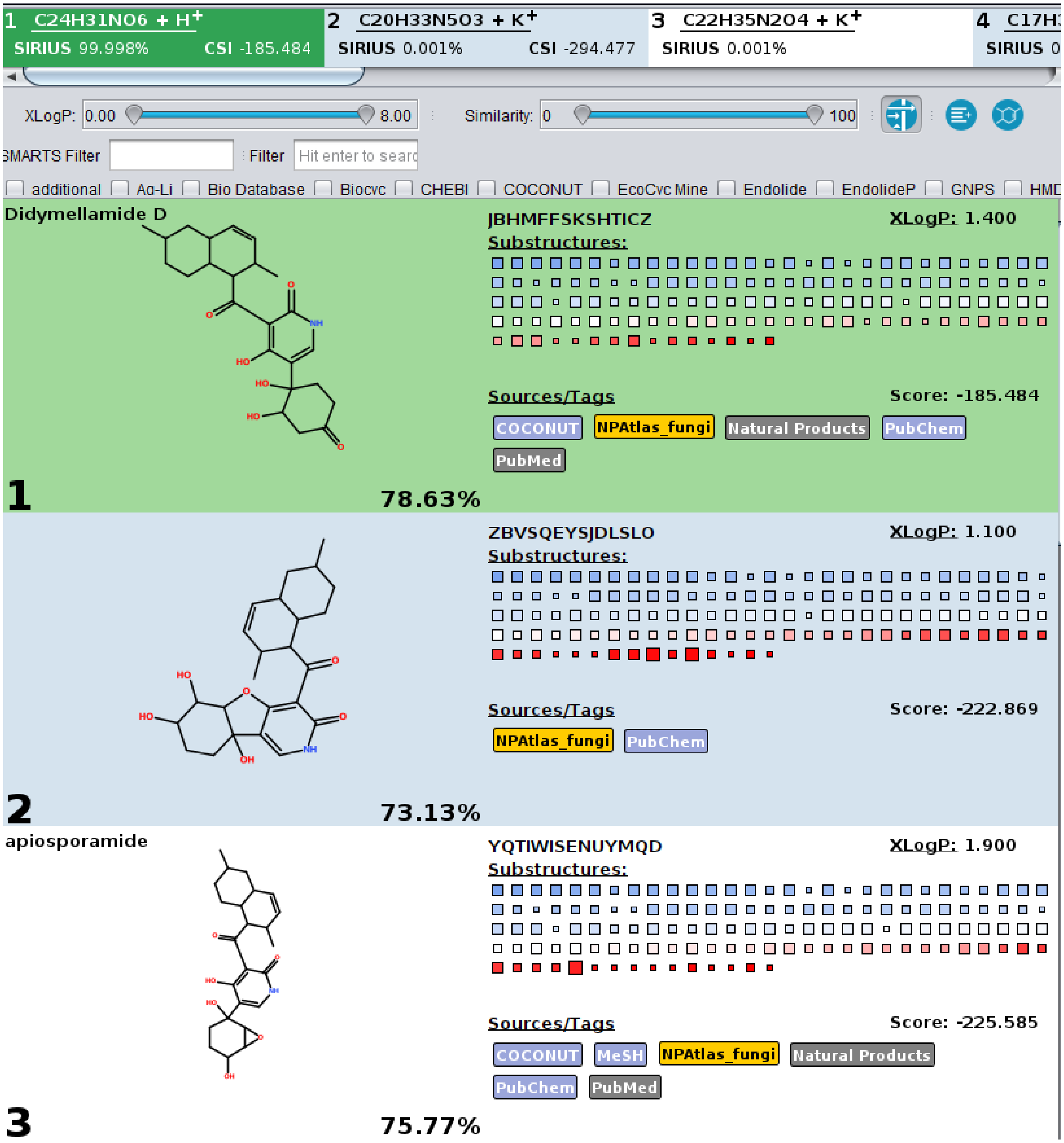

**Supplementary Figure 7: SIRIUS[2] molecular formula and CSI:FingerID[3] structure assumptions for AP.**

| $^{1}H$ | $^{13}C$ | Intensity |
|---|---|---|
| 8.00 | 133.0 | 861.8 |
| 5.59 | 130.9 | 1805.77 |
| 5.44 | 131.0 | 1864.71 |
| 4.31 | 53.4 | 972.04 |
| 4.22 | 63.7 | 839.31 |
| 3.57 | 56.2 | 20.01 |
| 3.57 | 58.7 | 447.89 |
| 2.81 | 31.2 | 633.72 |
| 2.00 | 25.8 | -12.73 |
| 1.9 | 28.5 | -17.05 |
| 1.86 | 28.5 | -12.77 |
| 1.86 | 29.9 | -10.2 |
| 1.75 | 35.4 | -11.57 |
| 1.75 | 41.7 | -423.11 |
| 1.60 | 36.0 | 735.44 |
| 1.52 | 25.8 | -10.15 |
| 1.48 | 33.0 | 420.28 |
| 1.04 | 35.4 | -16.29 |
| 0.91 | 22.6 | 11.07 |
| 0.88 | 29.9 | -5.63 |
| 0.85 | 18.1 | 6.49 |

**Supplementary Table 1: $^{1}H$-$^{13}C$-HSQC data table of NHAP used for SMART analysis.[4]**

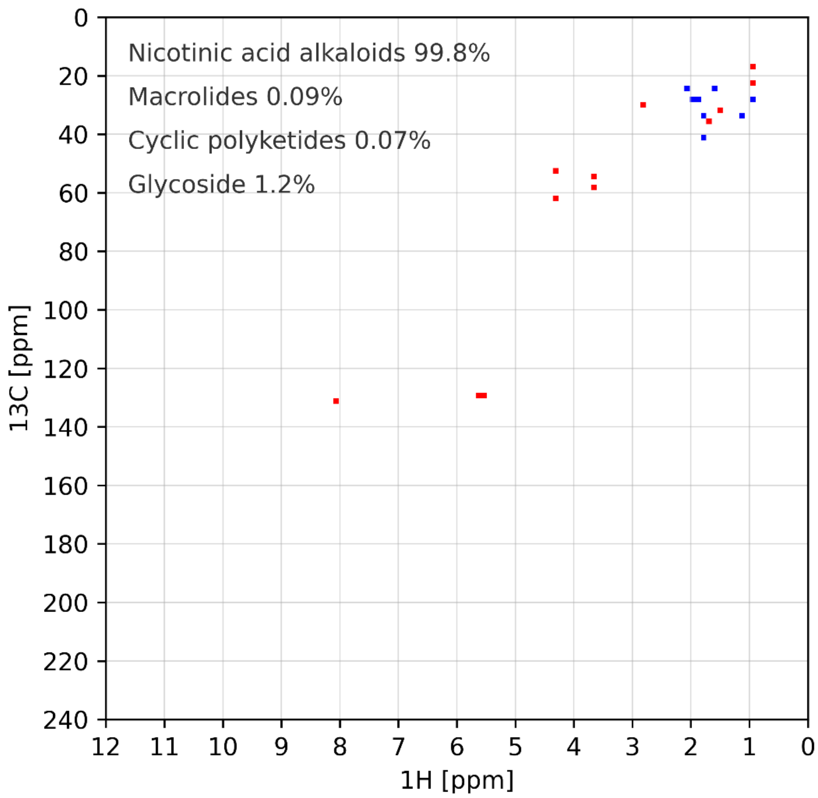

**Supplementary Figure 8: SMART compound class predictions for NHAP.[4]**

| | | |
|---|---|---|
| **1** | *N*-hydroxyapiosporamide (**1**) cosine score: 0.94 | 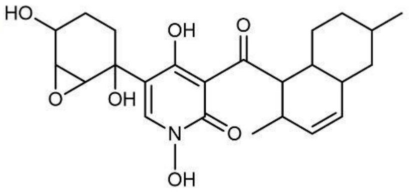 |
| **2** | apiosporamide (**2**) cosine score: 0.90 | 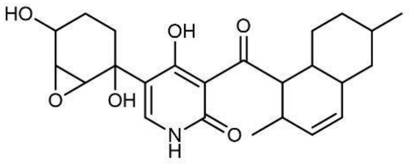 |
| **3** | arthpyrone C cosine score: 0.89 | 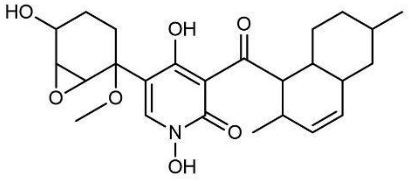 |

**Supplementary Figure 9: Top 3 SMART predictions for NHAP.[4]**

| $^{1}H$ | $^{13}C$ | Intensity |
|---|---|---|
| 7.83 | 138.9 | -1395.49 |
| 5.54 | 131.4 | -3317.21 |
| 5.39 | 131.2 | -3060.27 |
| 4.29 | 63.7 | -1290.69 |
| 4.29 | 53.1 | -66.22 |
| 3.56 | 55.4 | -984.85 |
| 3.47 | 57.8 | -981.18 |
| 2.82 | 31.4 | -2627.54 |
| 2.00 | 26.2 | 1227.35 |
| 1.90 | 28.8 | 59.75 |
| 1.85 | 30.0 | 67.05 |
| 1.80 | 42.0 | -86.49 |
| 1.80 | 28.8 | 65.56 |
| 1.73 | 35.4 | 77.98 |
| 1.73 | 41.8 | 67.83 |
| 1.59 | 36.3 | -2145.25 |
| 1.50 | 26.1 | 56.42 |
| 1.47 | 33.3 | -2208.76 |
| 1.04 | 35.4 | 36.70 |
| 0.90 | 30.0 | 6.78 |
| 0.90 | 22.7 | -1610.91 |

**Supplementary Table 2: $^{1}H$-$^{13}C$-HSQC data table of AP used for SMART analysis.**[4]

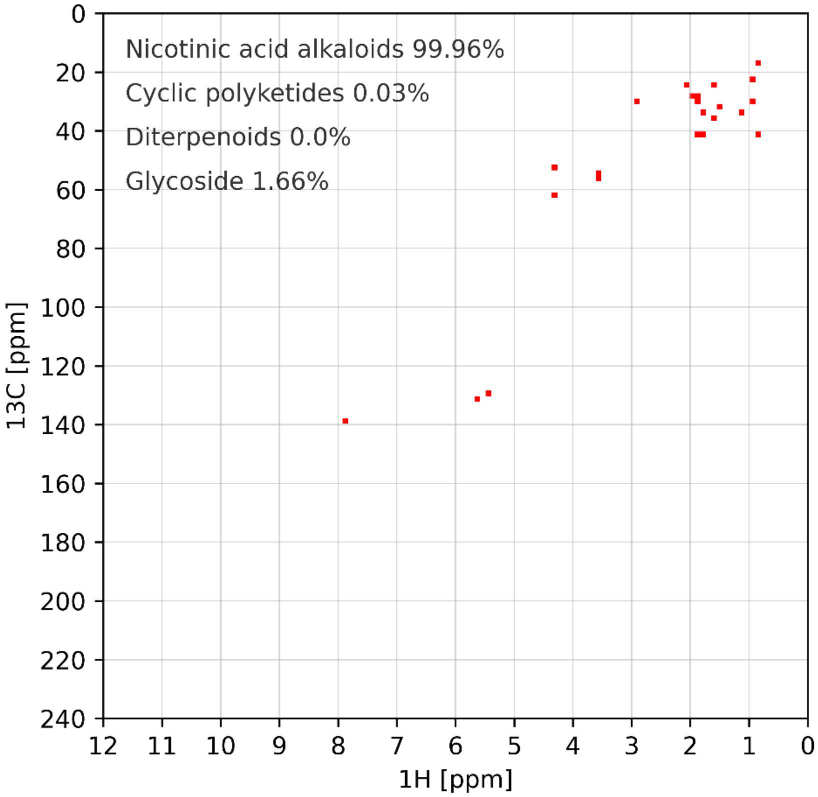

**Supplementary Figure 10: SMART compound class predictions for AP.**[4]

| | | |
|---|---|---|
| **1** | apiosporamide (**2**) cosine score: 0.95 | 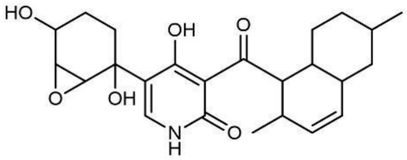 |
| **2** | *N*-hydroxyapiosporamide (**1**) cosine score: 0.93 | 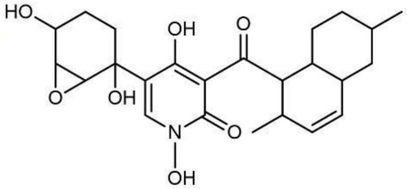 |
| **3** | arthpyrone C cosine score: 0.91 | 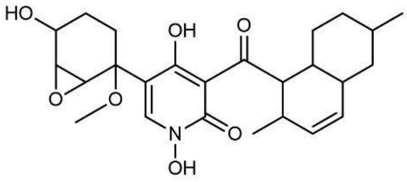 |

**Supplementary Figure 11: Top 3 SMART predictions for AP.**[4]

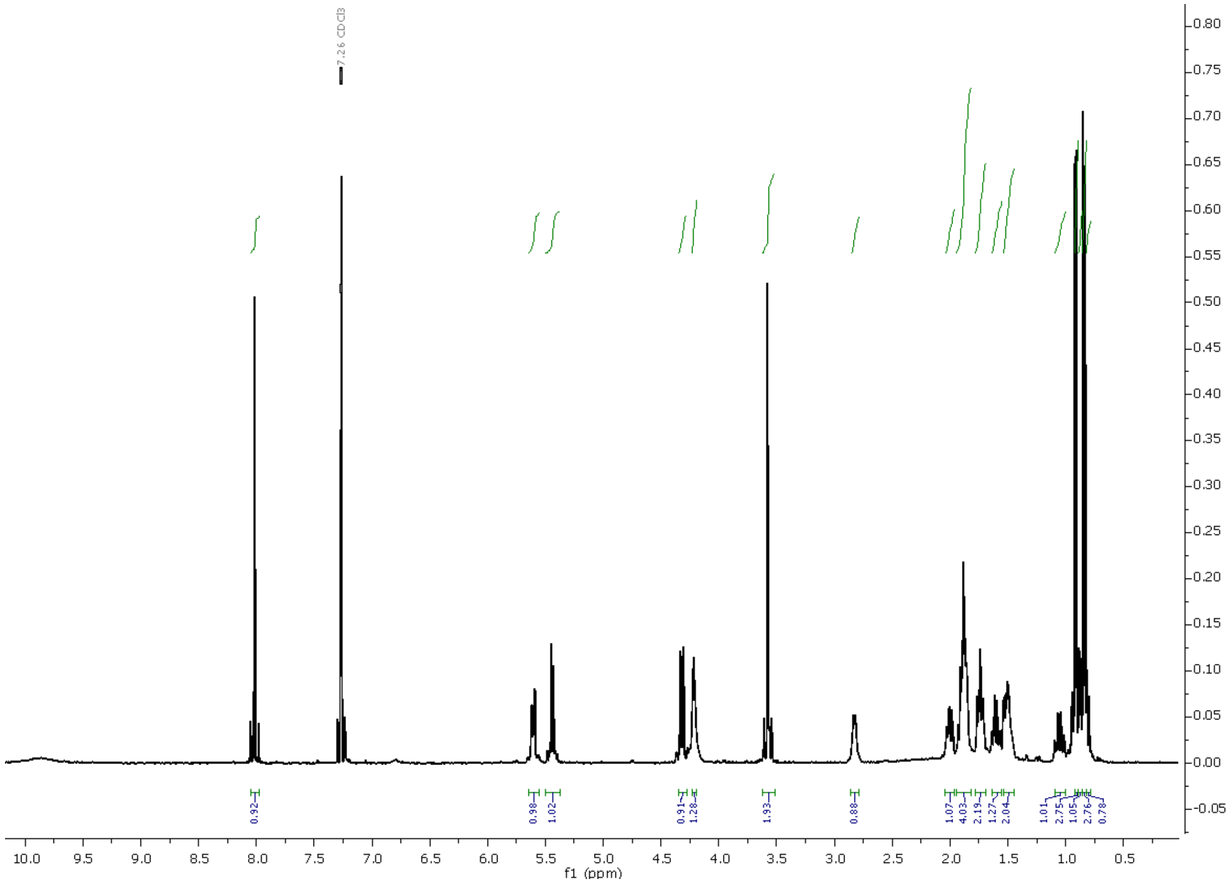

**Supplementary Figure 12: $^1$H NMR spectrum (500 MHz, chloroform-$d_1$) of NHAP.**

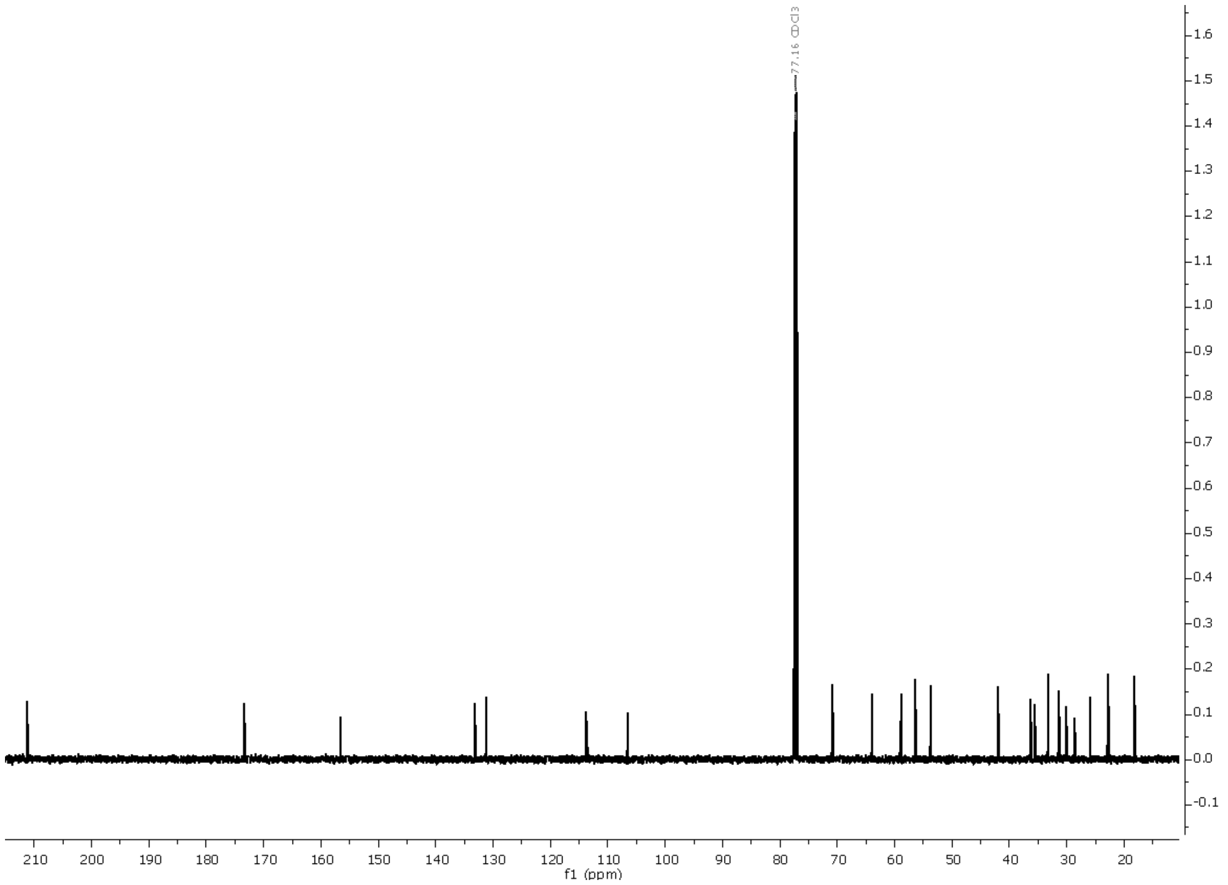

**Supplementary Figure 13: $^{13}$C NMR spectrum (125 MHz, chloroform-*$d_1$*) of NHAP.**

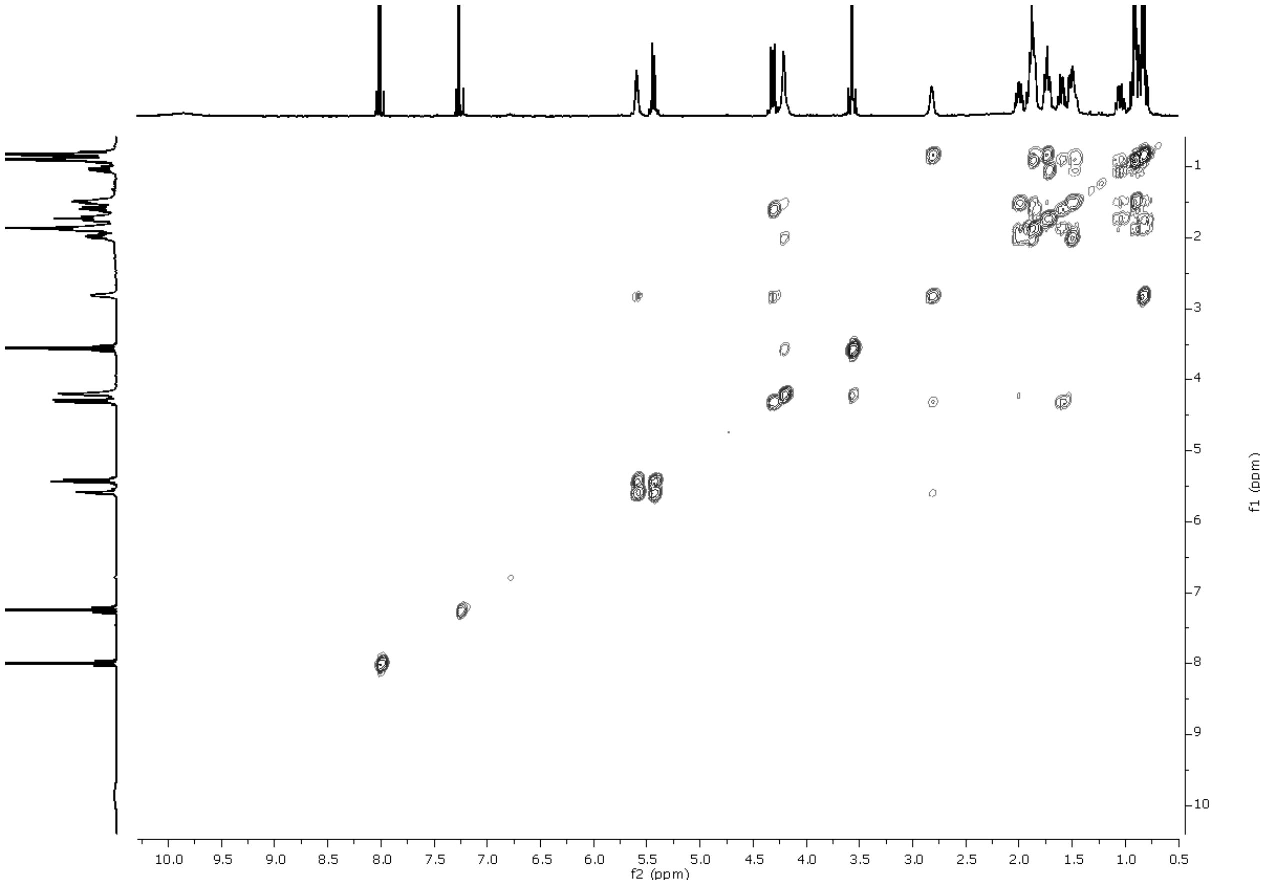

**Supplementary Figure 14: $^{1}$H-$^{1}$H-COSY NMR spectrum (500 MHz, chloroform-$d_1$) of NHAP.**

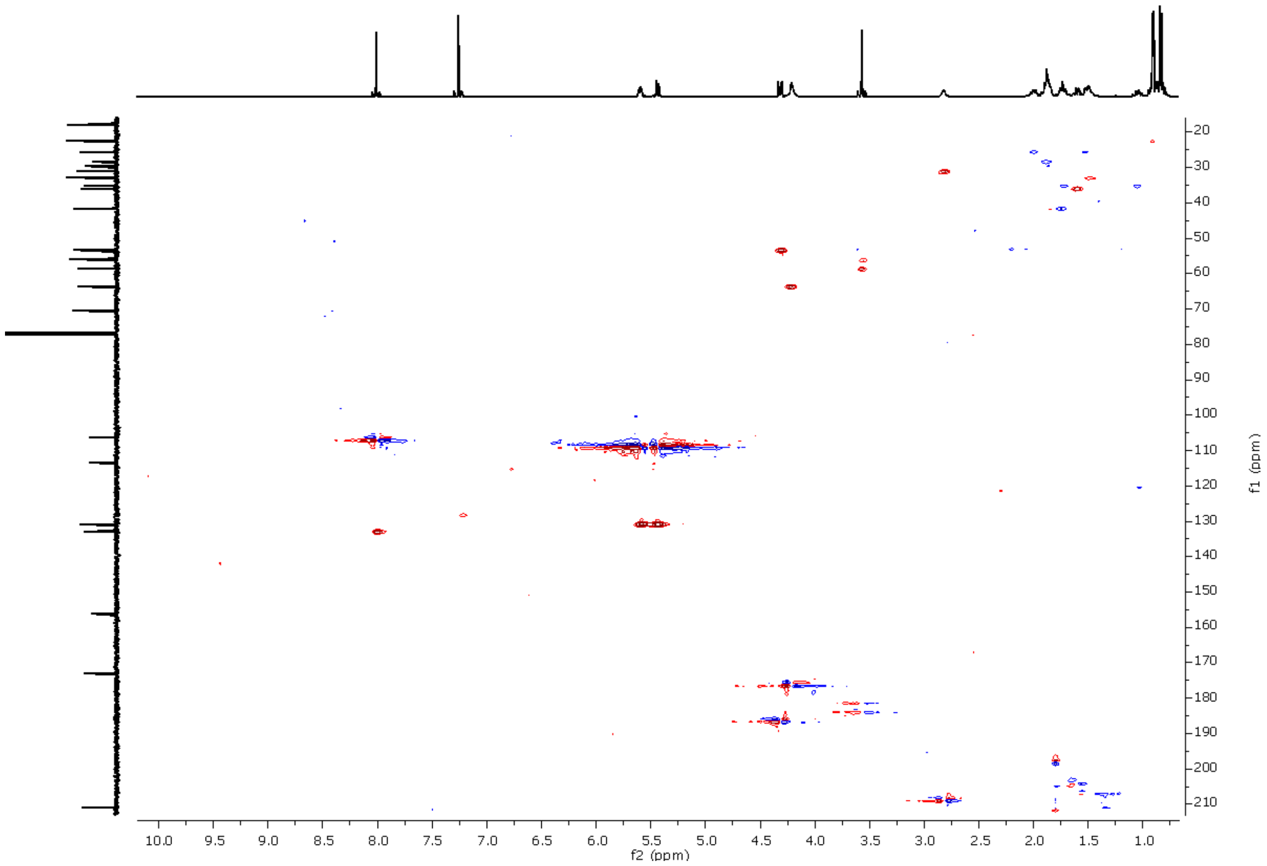

**Supplementary Figure 15: $^{1}H$-$^{13}C$-HSQC NMR spectrum (500 MHz, chloroform-*$d_1$*) of NHAP.**

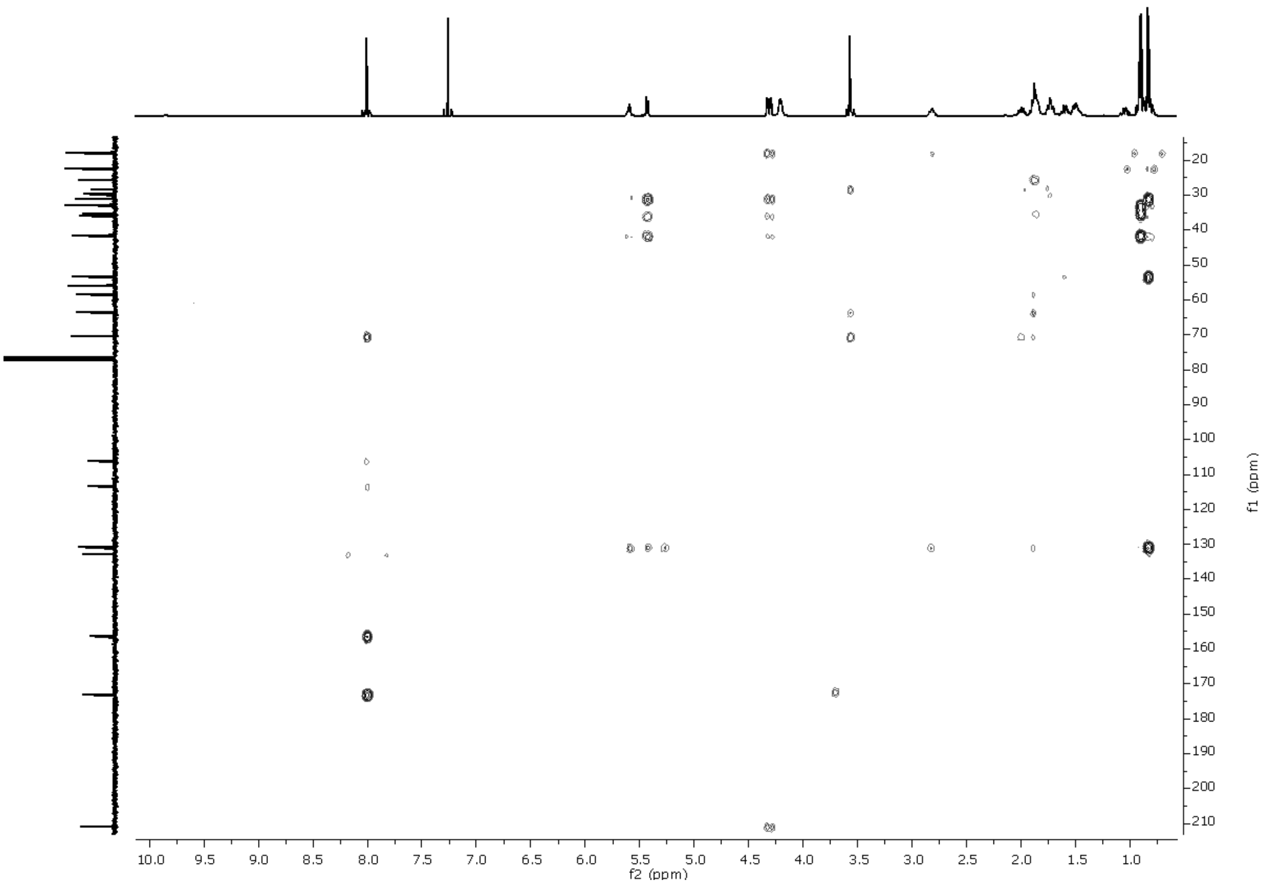

**Supplementary Figure 16: $^{1}H$-$^{13}C$-HMBC NMR spectrum (500 MHz, chloroform-$d_1$) of NHAP.**

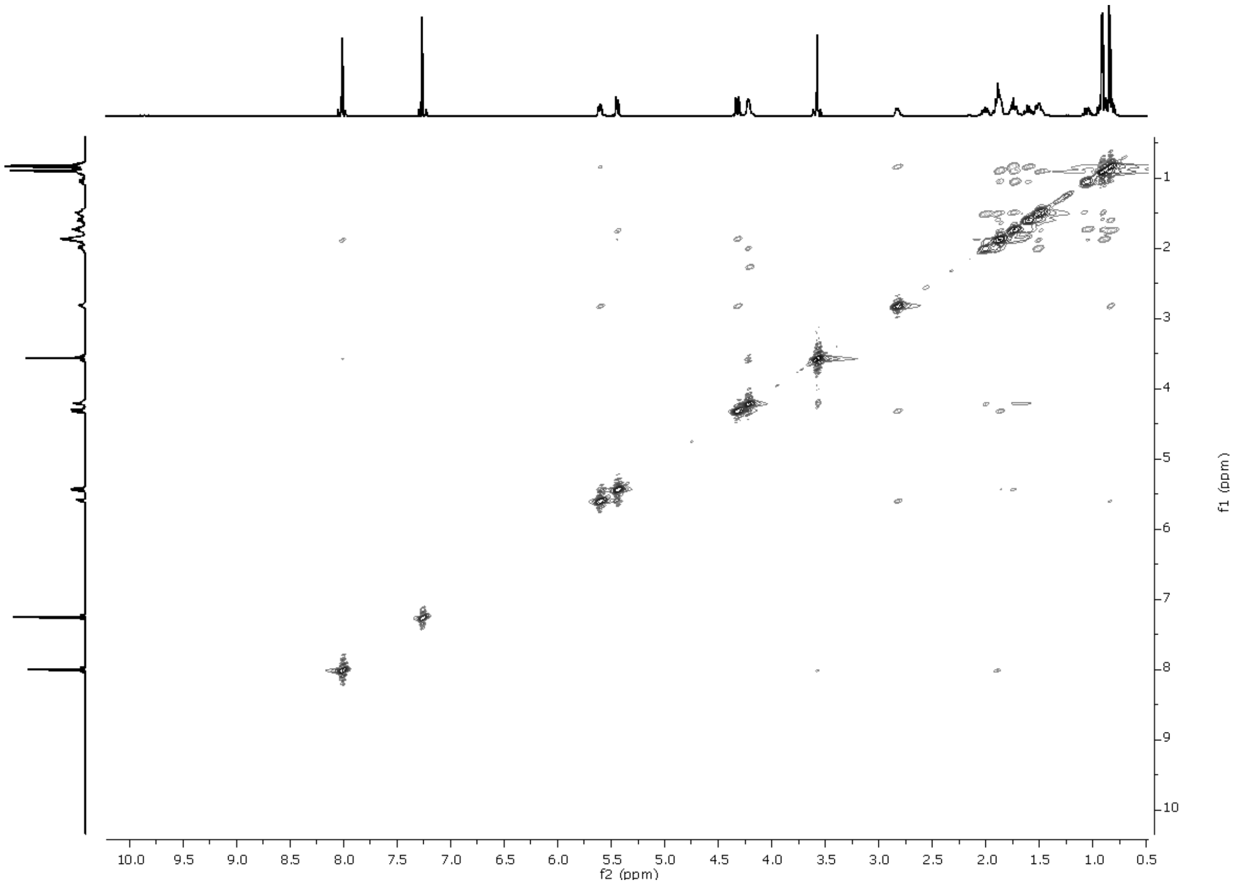

**Supplementary Figure 17: $^{1}H$-$^{1}H$-NOESY NMR spectrum (500 MHz, chloroform-$d_1$) of NHAP.**

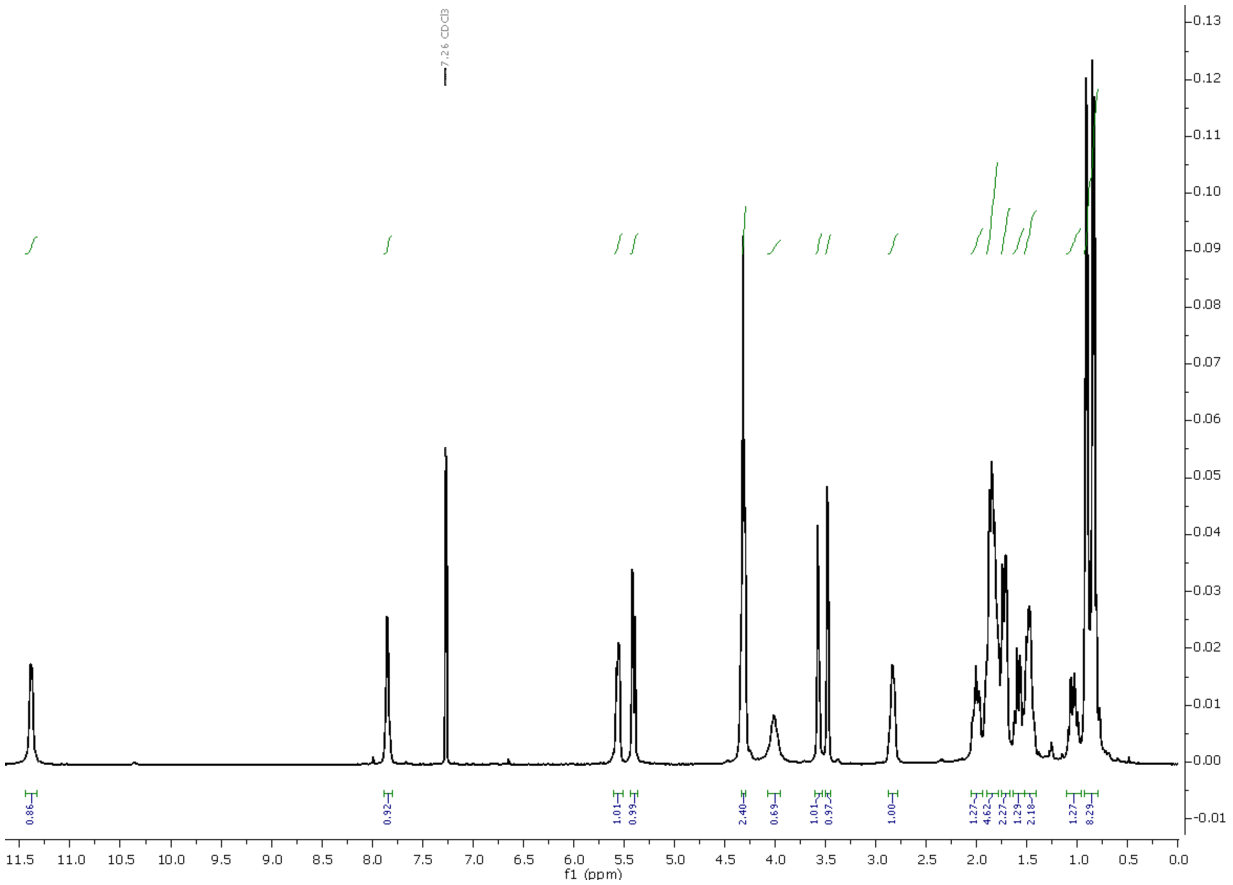

**Supplementary Figure 18: $^{1}$H NMR spectrum (400 MHz, chloroform-$d_1$) of AP.**

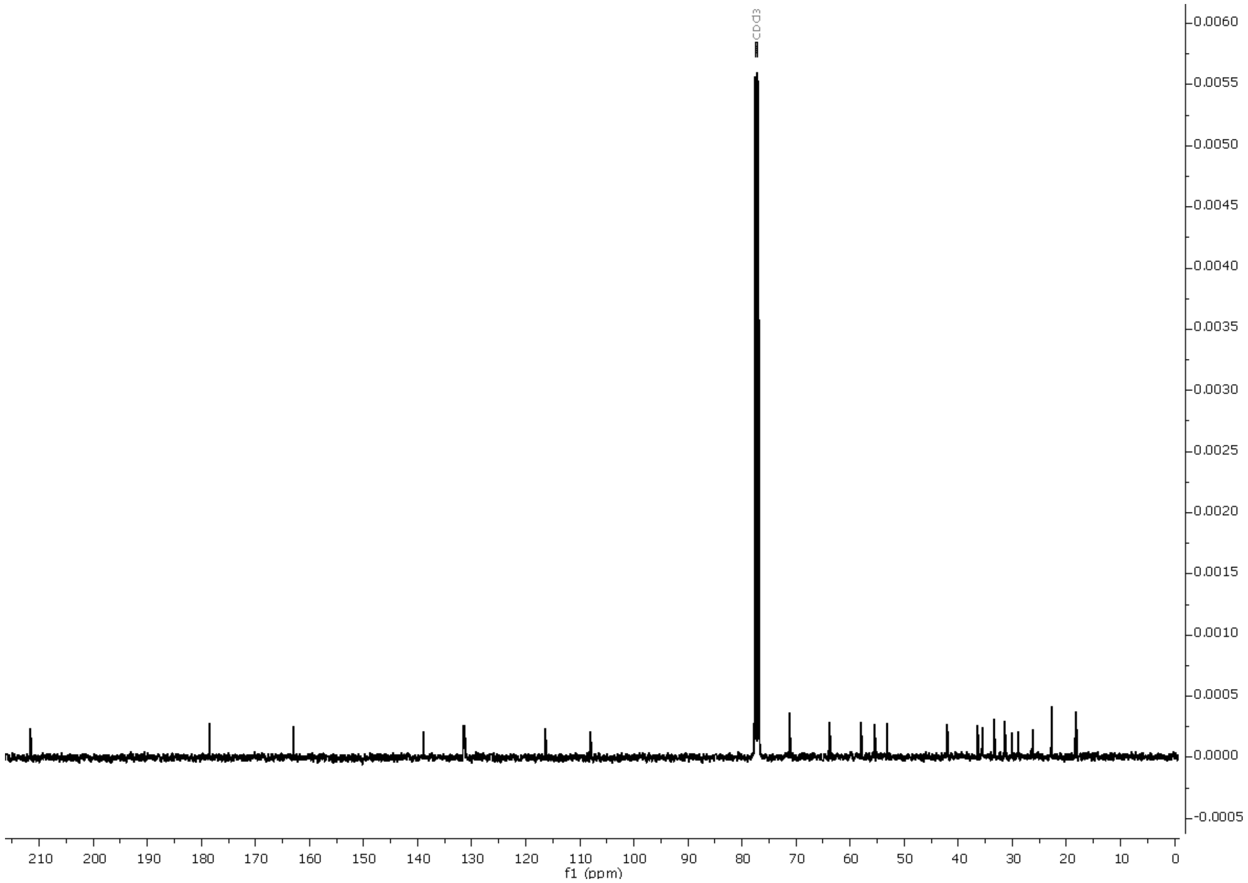

**Supplementary Figure 19: $^{13}$C NMR spectrum (100 MHz, chloroform-*$d_1$*) of AP.**

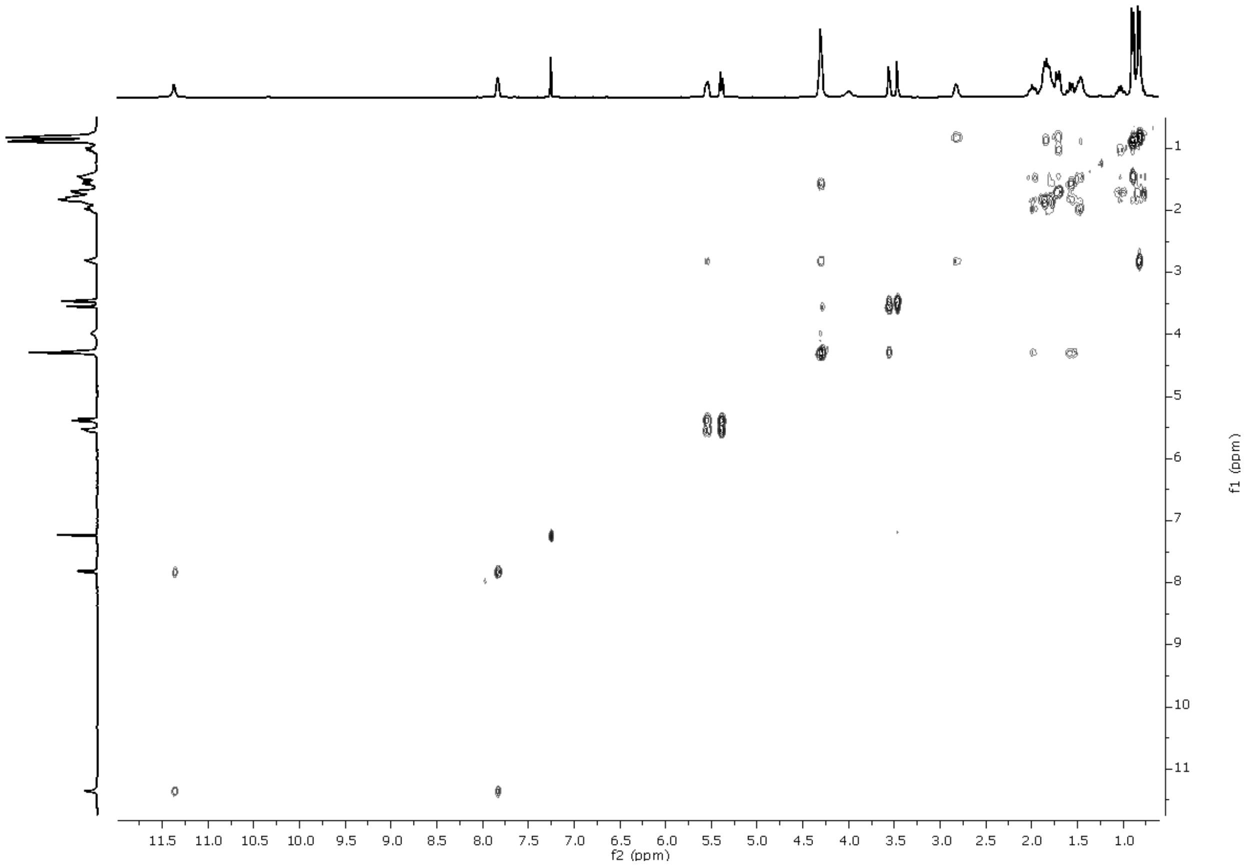

**Supplementary Figure 20: $^{1}H$-$^{1}H$-COSY spectrum (400 MHz, chloroform-$d_1$) of AP.**

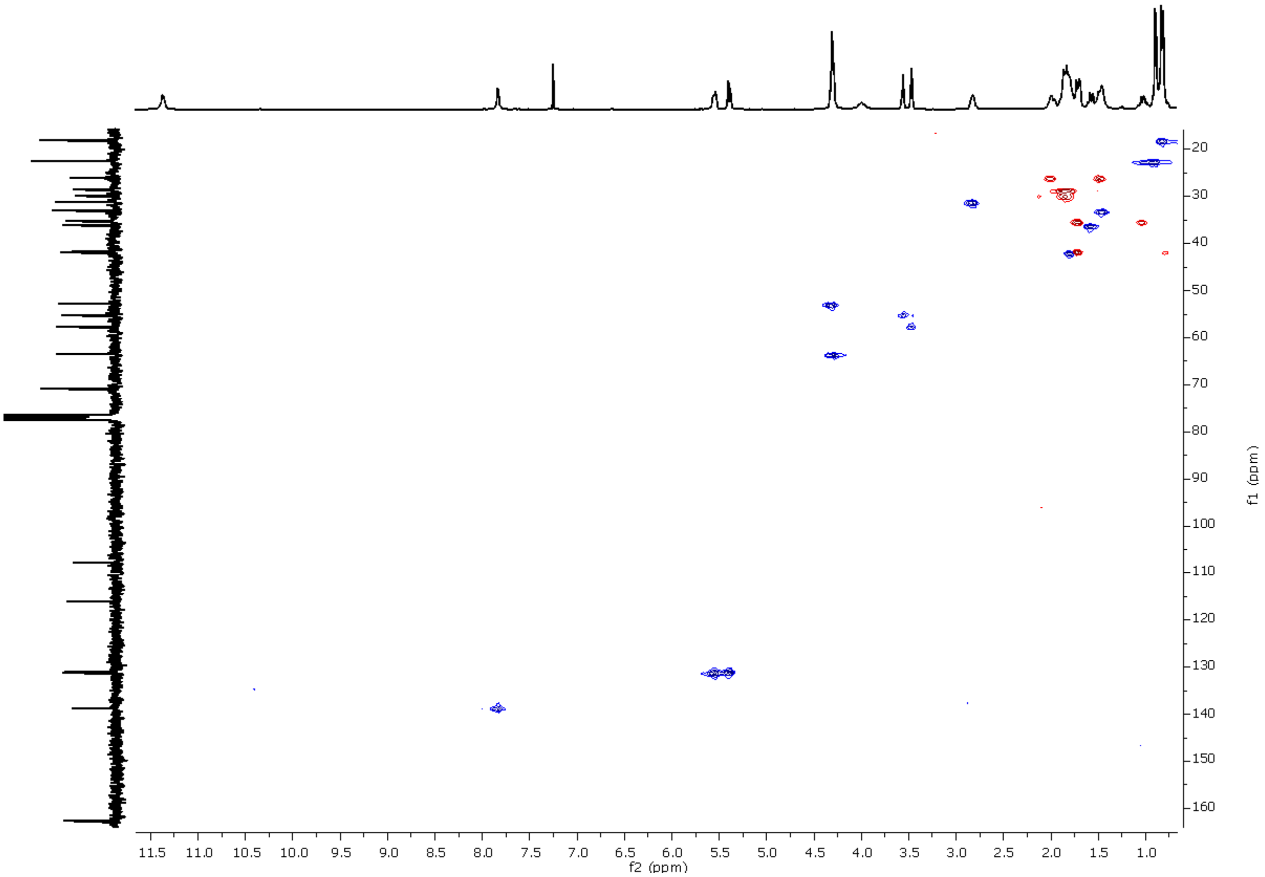

**Supplementary Figure 21: $^{1}H$-$^{13}C$-HSQC spectrum (400 MHz, chloroform-$d_1$) of AP.**

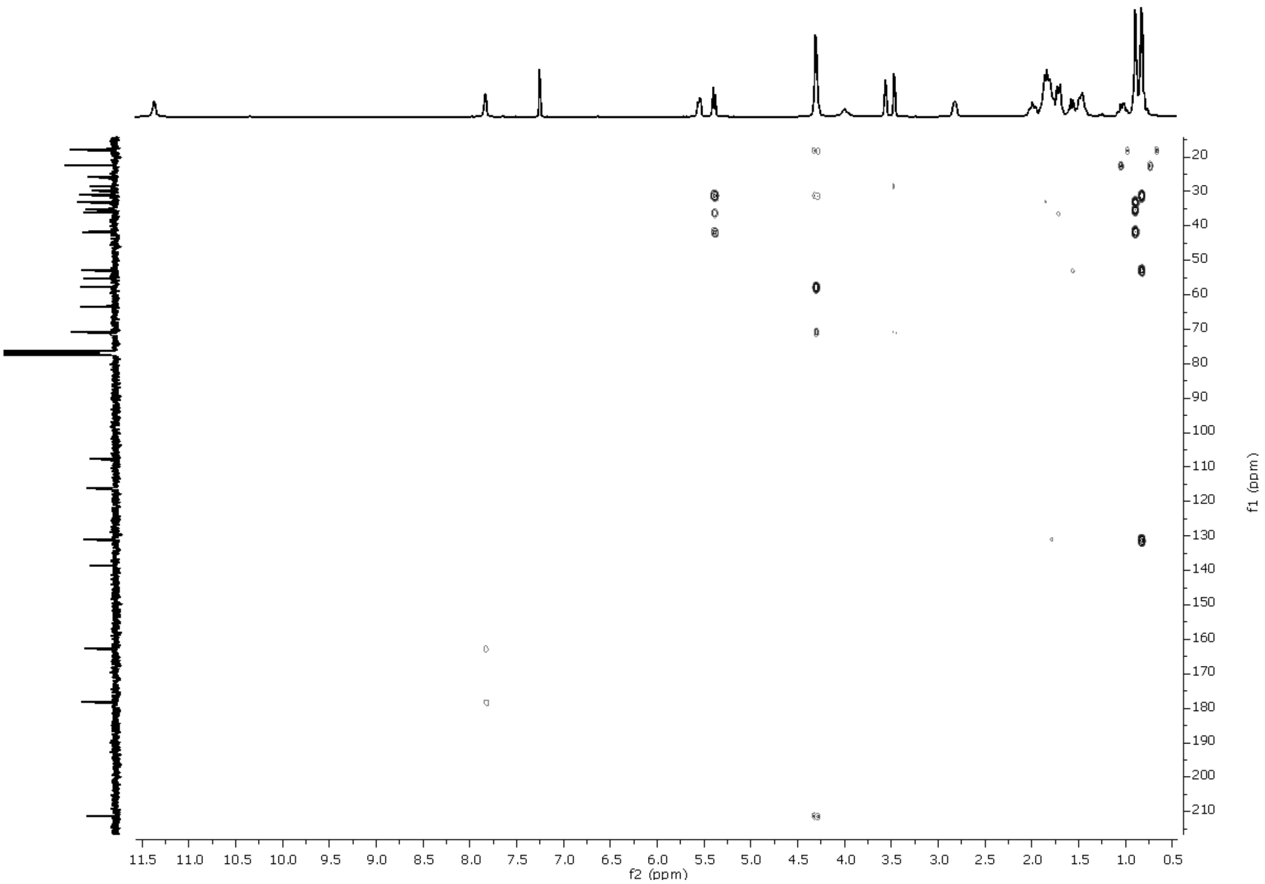

**Supplementary Figure 22: $^{1}$H-$^{13}$C-HMBC spectrum (400 MHz, chloroform-$d_1$) of AP.**

| | *N*-hydroxyapiosporamide (reference) $CD_3OD$ | | *N*-hydroxyapiosporamide (experimental) $CDCl_3$ | |
|---|---|---|---|---|
| **position** | **δC, type** | **δH (*J* in Hz)** | **δC, type** | **δH (*J* in Hz)** |
| 1 | - | - | - | - |
| 2 | 159.6,C | - | 156.5, C | - |
| 3 | 108.6, C | - | 106.4, C | - |
| 4 | 175.6, C | - | 173.2, C | - |
| 5 | 114.6, C | - | 113.7, C | - |
| 6 | 140.0, CH | 8.03 s | 133.1, CH | 8.01 s |
| 7 | 211.7, C | - | 211.1, C | - |
| 8 | 54.5, CH | 4.43 dd (5.2, 11.1) | 53.6, CH | 4.31 dd (5.6, 11.3) |
| 9 | 37.6, CH | 1.56 m | 36.2, CH | 1.60 m |
| 10 | 30.9, $CH_2$ | 0.88 m<br>1.92 m | 29.9, $CH_2$ | 0.87 m<br>1.89 m |
| 11 | 36.6, $CH_2$ | 0.88 m<br>1.92 m | 35.4, $CH_2$ | 1.05 m<br>1.72 m |
| 12 | 34.3, CH | 1.49 m | 33.2, CH | 1.48 m |
| 13 | 43.1, $CH_2$ | 0.77 m<br>1.73 m | 41.8, $CH_2$ | 0.80 m<br>1.75 m |
| 14 | 43.2, CH | 1.81 m | 41.8, CH | 1.85 m |
| 15 | 131.7, CH | 5.40 brd (9.8) | 131.1, CH | 5.44 brd (10.0) |
| 16 | 132.5, CH | 5.59 m | 131.1, CH | 5.60 m |
| 17 | 32.4, CH | 2.84 s | 31.3, CH | 2.82 m |
| 18 | 18.4, $CH_3$ | 0.81 d (6.9) | 18.1, $CH_3$ | 0.84 d (7.1) |
| 19 | 22.9, $CH_3$ | 0.92 d (6.5) | 22.6, $CH_3$ | 0.91 d (6.8) |
| 20 | 70.4, C | - | 70.7, C | - |
| 21 | 60.5, CH | 3.63 brs | 58.8, CH | 3.57 brs |
| 22 | 57.1, CH | 3.41 brs | 56.3, CH | 3.57 brs |
| 23 | 67.1, CH | 4.12 m | 63.8, CH | 4.21 m |
| 24 | 25.7, $CH_2$ | 1.36 m<br>1.80 m | 25.8, $CH_2$ | 1.52 m<br>2.00 m |
| 25 | 31.7, $CH_2$ | 1.67 m<br>2.24 m | 28.5, $CH_2$ | 1.87 m<br>1.91 m |

**Supplementary Table 3: NMR Spectroscopic Data (500 MHz) in methanol-*$d_4$* for *N*-hydroxyapiosporamide[5] and (400MHz) *N*-hydroxyapiosporamide (isolated in this study) in chloroform-*$d_1$*.**

| apiosporamide (reference) acetone-$d_6$ | | | apiosporamide isolated (experimental) $CDCl_3$ | |
|---|---|---|---|---|
| position | δC, type | δH (*J* in Hz) | δC, type | δH (*J* in Hz) |
| 1 | - | 10.46 s | - | 11.38 d (4.2) |
| 2 | 162.2, C | - | 162.9, C | - |
| 3 | 108.4, C | - | 107.9, C | - |
| 4 | 179.9, C | OH 11.57 s | 178.4, C | - |
| 5 | 115.9, C | - | 116.2, C | - |
| 6 | 140.0, CH | 7.66 s | 138.9, CH | 7.84 d (4.2) |
| 7 | 211.4, C | - | 211.5, C | |
| 8 | 54.2, CH | 4.45 dd | 53.0, CH | 4.30 m |
| 9 | 37.7, CH | 1.68 m | 36.3, CH | 1.58 m |
| 10 | 31.0, $CH_2$ | 0.90 m | 30.0, $CH_2$ | 0.86 m |
| | | 1.93 m | | 1.87 m |
| 11 | 36.2, $CH_2$ | 0.99 m | 35.4, $CH_2$ | 1.04 m |
| | | 1.76 m | | 1.72 m |
| 12 | 33.8, CH | 1.56 m | 33.2, CH | 1.47 m |
| 13 | 42.6, $CH_2$ | 0.74 m | 41.8, $CH_2$ | 0.81 m |
| | | 1.76 m | | 1.72 m |
| 14 | 42.6, CH | 1.87 m | 42.0, CH | 1.81 m |
| 15 | 131.3, CH | 5.39 | 131.1, CH | 5.40 d (9.9) |
| 16 | 132.5, CH | 5.59 m | 131.3, CH | 5.56 m |
| 17 | 31.8, CH | 2.84 s | 31.3, CH | 2.83 m |
| 18 | 18.3, $CH_3$ | 0.81 | 18.2, $CH_3$ | 0.83 d (7.0) |
| 19 | 22.8, $CH_3$ | 0.90 | 22.7, $CH_3$ | 0.90 d (6.4) |
| 20 | 70.2, C | OH 4.35 | 71.0, C | - |
| 21 | 60.0, CH | 3.54 brs | 55.4, CH | 3.48 d (3.5) |
| 22 | 57.3, CH | 3.34 brs | 57.9, CH | 3.57 t (3.6) |
| 23 | 66.4, CH | 4.08 m, OH 3.83 | 63.6, CH | 4.31 m |
| 24 | 26.0, $CH_2$ | 1.33 m | 26.1, $CH_2$ | 1.49 m |
| | | 1.87 m | | 2.00 m |
| 25 | 31.5, $CH_2$ | 1.71 m | 28.8, $CH_2$ | 1.81 m |
| | | 2.17 m | | 1.90 m |

**Supplementary Table 4: NMR Spectroscopic Data (300 MHz) in acetone-$d_6$ for apiosporamide[6] and (400MHz) apiosporamide (isolated in this study) in chloroform-$d_1$.**

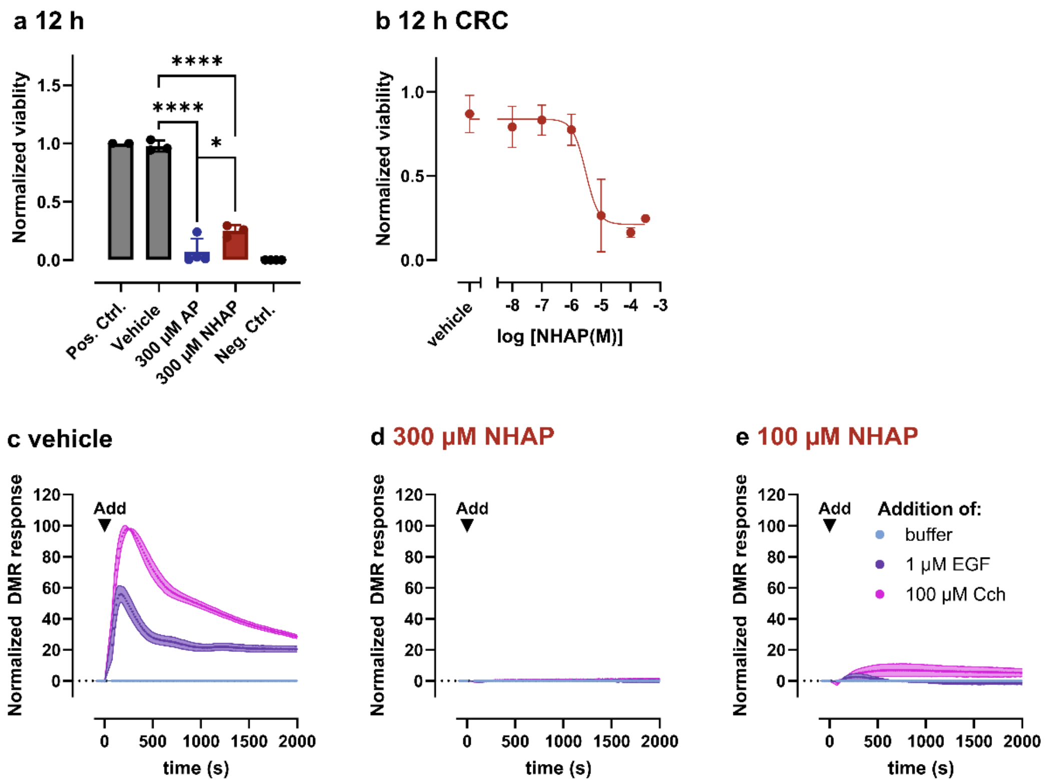

**Supplementary Figure 23: Effects of NHAP and AP on HEK293 cell viability and signaling responses.**
Cell viability and cytotoxicity were assessed using CellTiter-Blue® assays and whole-cell biosensing based on detection of dynamic mass redistribution (DMR). a) Normalized CellTiter-Blue fluorescence (Ex/Em 560/590 nm) as a measure of cell viability. Cells were treated with the indicated concentrations of NHAP, AP, or vehicle for 12 h, followed by incubation with CellTiter-Blue reagent for 2 h. b) Concentration-response curve for NHAP. Data were normalized to untreated cells (100%) and medium-only controls (0%) and are shown as mean ± SEM of 3–4 independent experiments. Statistical analysis was performed using one-way ANOVA followed by multiple comparisons testing (****,p < 0.0001). c-e) Normalized DMR response (expressed as % of carbachol (Cch)-induced amplitude) in HEK293 cells pre-incubated for 12 h with c) vehicle, d) 300 µM NHAP, or e) 100 µM NHAP. Cells were subsequently stimulated via endogenous epidermal growth factor receptor (EGFR) signaling using EGF or muscarinic acetylcholine M3 receptor signaling using Cch. Data are shown as mean ± SEM of 2–3 independent experiments.

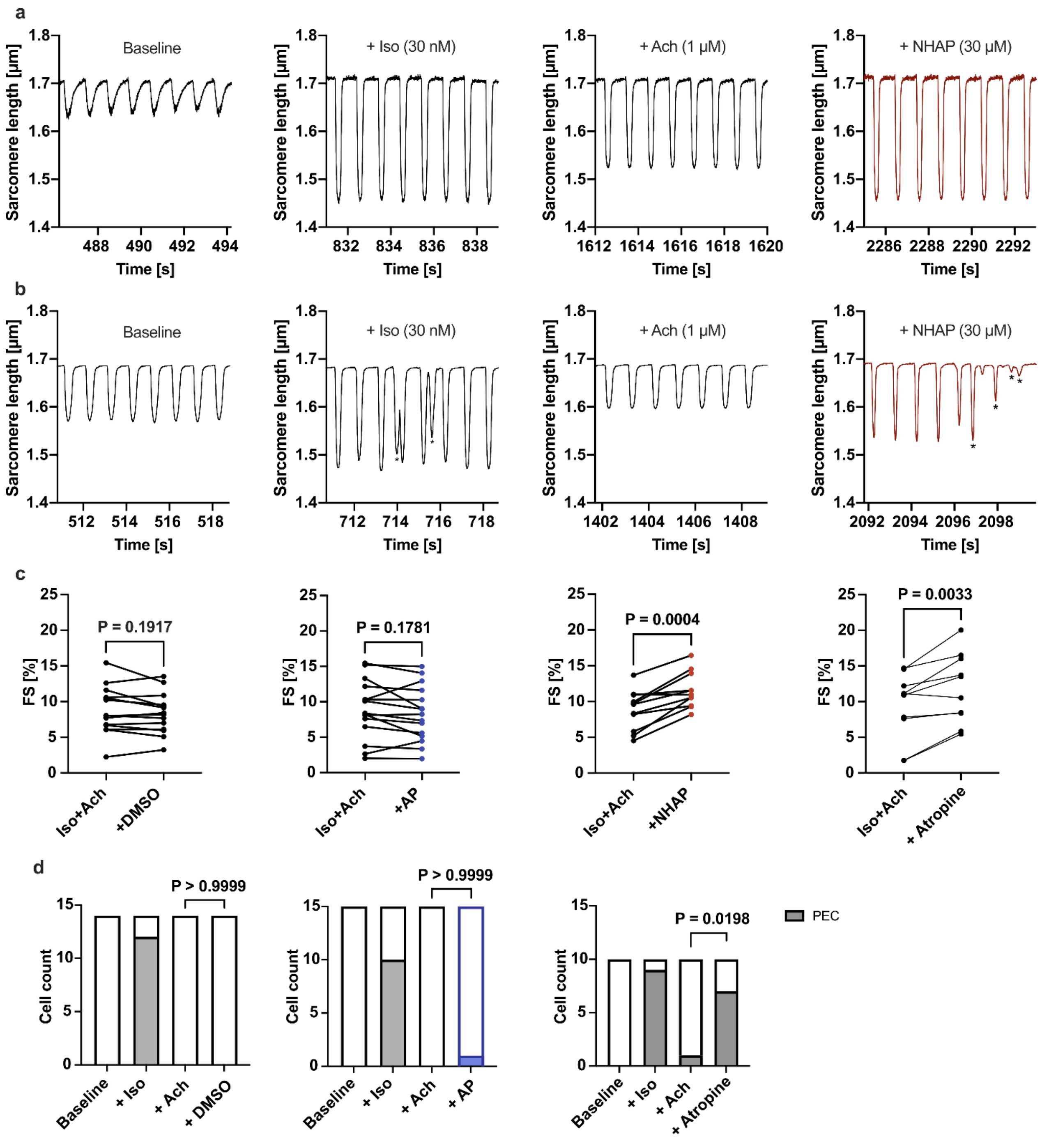
a
Baseline
+ Iso (30 nM)
+ Ach (1 µM)
+ NHAP (30 µM)
Sarcomere length [µm]
Time [s]
b
Baseline
+ Iso (30 nM)
+ Ach (1 µM)
+ NHAP (30 µM)
Sarcomere length [µm]
Time [s]
c
FS [%]
P = 0.1917
Iso+Ach
+DMSO
P = 0.1781
Iso+Ach
+AP
P = 0.0004
Iso+Ach
+NHAP
P = 0.0033
Iso+Ach
+ Atropine
d
Cell count
P > 0.9999
Baseline
+ Iso
+ Ach
+ DMSO
P > 0.9999
Baseline
+ Iso
+ Ach
+ AP
P = 0.0198
Baseline
+ Iso
+ Ach
+ Atropine
PEC

**Supplementary Figure 24: Effects of NHAP, AP, and atropine on rat ventricular cardiomyocytes.**

**a** Individual sarcomere shortening recordings of a cardiomyocyte. Isoprenaline (Iso) increases sarcomere shortenings compared to baseline conditions. The following negative-inotropic effect of acetylcholine (Ach) is reversed by NHAP. **b** Individual sarcomere shortening recordings of the cardiomyocyte shown in Figure 5b (lower left panel). Here, Iso elicits pro-arrhythmic extra contractions (PEC, marked by asterisks) in addition to its positive-inotropic effect. Ach suppresses the PEC and causes a negative-inotropic effect. Finally, NHAP leads to the re-occurrence of the PEC and elicits a positive-inotropic effect. **c** Time matched comparison of fractional shortening (FS) before and after the addition of DMSO, AP, NHAP or Atropine in each cell (Paired t-test). **d** Occurrence of PEC at baseline conditions and following the addition of Iso (+ Iso), Ach (+ Ach), and either DMSO (+ DMSO), AP (+ AP) or Atropine (+ Atropine). In contrast to NHAP (see Figure 5d), neither DMSO nor AP led to the re-occurrence of PEC. Atropine had comparable effects on PEC as NHAP (Fisher’s exact test). Data were obtained from the following numbers of animals (N) and cells (n) per group: DMSO: N=7, n=14; AP: N=5, n=15; NHAP: N=5, n=11; Atropine: N=5; n=10.

### Methods:

#### Proof-of-concept:

Purified G$\alpha_{i1}$, G$\alpha_s$, G11-heterotrimer, and FR-sensitive G$\alpha_{i1}$ were buffer exchanged in 100 mM ammonium acetate (pH 7.5) and incubated with FR900359 prior to native MS measurements. All native MS measurements were conducted with a collision energy of 10.

#### Selectivity test with endolide E:

Purified G$\alpha_{i1}$, G$\alpha_s$, G11-heterotrimer were buffer exchanged in 100 mM ammonium acetate prior to native MS measurements. Endolide E was analyzed at a concentration ten times higher than that of the respective G-protein.

All native MS analyses were carried out using a spray voltage of 3 kV with S-lens RF level at 70. The probe heater temperature was set to 50 °C and heated capillary temperature to 253°C. The sheath gas flow rate was set to 20, the aux gas flow rate to 7 and the sweep gas flow rate to 2. A higher collision energy of 10 was conducted, respectively.

A solution of 40 µM G$\alpha_{i1}$ is incubated with an equal volume of 7 M urea.[7] The urea is then exchanged using 100 mM ammonium acetate. After buffer exchange an excess of 10-fold endolide E is added prior to the native MS measurement.

#### Structure elucidation of NHAP/AP:

The key correlations in NHAP are firstly the COSY-spin system which occurs from H-8, H-9, H-10, and H-14 to H-18. Together with the HMBC correlations from H-19 to C-13 and H-8 to C-18, it reveals a decalin moiety that is twice-methylated at the 12$^{th}$ and 17$^{th}$ positions. Second, the HMBC correlation of H-8 to C-7 shows the bond of the decalin moiety to the 4-hydroxy-2-pyridone derivative.

Further HMBC correlations from H-6 to C-2, C-4, and C-20 with the NOESY correlation from H-6 to H-25 show the linkage to the epoxycyclohexane-diol moiety. Within this scaffold, in addition to the HMBC correlations from H-25 to C-21 and C-23, a COSY-spin system from H-22 to H-24 can be observed.

Analogous correlations can be found in AP. The decalin moiety also has a COSY-spin system from H-14 to H-15 via H-9 with H-18 and from H-10 to H-13 with H-19. Additionally, there is a HMBC correlation between H-8 and C-7. Furthermore, the epoxycyclohexane-diol moiety has a COSY-spin system from H-21 to H-25, so that the general structure follows that of NHAP. Considering the Δ*m/z* of 15.9942, there is evidence for a structural difference of one oxygen atom. Observation of the signal $\delta_H$ 11.38 only in the $^1$H NMR spectrum of AP, indicating an *N*-H signal (**Fig. S12**). The lack of the *N*-hydroxy group in AP is clearly confirmed.

The relative structure of NHAP can also be determined from the most significant NOESY correlations. In the epoxycyclohexane-diol moiety there are NOESY correlations between H-23 to H-21 and to H-22. These hydrogens therefore have the same configuration. Previously, decalin biosynthesis was shown to be possible in the trans configuration for NHAP and in the cis configuration for Fischerin, a closely related compound to NHAP.[8] The arrangement of the ring system in the investigated compounds can be determined to be trans, since there is no NOESY correlation between H-9 and H-14, nor between H-9 and H-8. In addition, H-8 and H-14 show NOESY correlations. Furthermore, H-17 shows a NOESY correlation with H-8, indicating that the C-18 methyl group is configured in the opposite direction. Epimers have also been reported for this dimethylated decalin moiety.[5] Further NOESY correlations from H-8 to H10-a support an opposite arrangement of 10-b, which shows correlations to 13-b, which further shows a correlation to H-12. Thus the C-19 methyl group has the same configuration as the C-18 methyl group. This strong evidence supports the previously proposed absolute structure of (-)-NHAP and (-)-AP.[6,9]
